# Age-Related Changes in Functional Balance Ability Predict Alpha Activity During Multisensory Integration

**DOI:** 10.1101/2024.12.17.628991

**Authors:** Jessica L. Pepper, Bo Yao, Jason J. Braithwaite, Theodoros M. Bampouras, Helen E. Nuttall

**Affiliations:** Department of Psychology Fylde College Lancaster University Lancaster, United Kingdom LA1 4YF; School of Sport and Exercise Sciences Liverpool John Moores University Liverpool, United Kingdom L3 3AF

**Keywords:** ageing, alpha, attention, balance, falls, multisensory

## Abstract

The increased multisensory integration and weaker attentional control experienced by older adults during audiovisual processing can result in inaccurate perceptions of their dynamic, everyday environment. These inaccurate representations of our environment can contribute to increased fall risk in older adults. A neural correlate of the attentional difference between younger and older adults could be oscillatory alpha activity (8-12Hz), indexing inhibitory processes during multisensory integration. The current study investigated whether age-related changes in alpha activity underlie weaker attentional control in older adults during a multisensory task, and if alpha associates with fall risk.

Thirty-six younger (18-35 years old) and thirty-six older (60-80 years old) adults completed a cued-spatial-attention stream-bounce task, assessing audiovisual integration when attending to validly-cued or invalidly-cued locations, at 0ms or 300ms stimulus-onset asynchronies. Oscillatory alpha activity was recorded throughout using EEG to index participants’ inhibitory abilities. Functional ability and balance were measured to index fall risk.

Multiple linear regression models revealed that even when attending to the validly-cued location, less accurate multisensory integration was exhibited by older adults compared to younger adults, suggesting that older adults demonstrate weaker top-down modulation of multisensory integration through failing to inhibit task-irrelevant information. However, alpha power across the trials did not predict the extent of multisensory integration within the task. A significant interaction between age and functional ability scores predicted alpha power, suggesting that older adults may rely on attentional mechanisms for functional ability more than younger adults do. Potential implications in the design of clinical treatments to reduce falls are discussed.

## Introduction

By 2050, it is expected that over 20% of the UK population will be 60 years old or above, with approximately 30% of community-dwelling adults over 65 suffering from falls (Zhang et al., 2020; Office for Health Improvement & Disparities, 2022). Falls have serious consequences on both an individual level and a systemic level – not only are they the most common cause of death for adults over 65, but also it is estimated that injuries associated with falls cost the National Health Service over £4.4 billion per year (Office for Health Improvement & Disparities, 2022). It is therefore highly important to understand the multifaceted causes of falls, including the age-related changes in perceptual and cognitive processes that may contribute to weaker functional ability and increased fall risk.

One potential reason behind increased fall risk in older adults are the age-related changes in multisensory integration. Multisensory integration describes the perceptual and cognitive mechanisms involved in binding sensory information together, to form a unitary percept of a person’s body and environment (Stevenson et al., 2012; Talsma et al., 2010; Stein & Wallace, 1996; Diederich & Colonius, 2004). Research suggests that older adults display increased multisensory integration relative to younger adults (Pepper et al., 2023; Pepper & Nuttall, 2023; Laurienti et al., 2006; Peiffer et al., 2007; Mahoney et al., 2011).

This increased integration has a positive outcome when the sensory information is congruent and should be integrated. For example, effectively utilising visual and auditory cues improves driving performance (Ramkhalawansingh et al., 2016) and speech perception abilities (see Jones & Noppeney, 2021, for a review). On the other hand, when task-irrelevant or incongruent information is erroneously integrated, this can have a negative outcome, producing representations of the environment that are confusing, noisy, and unstable (de Dieuleveult et al., 2017; Bedard & Barnett-Cowan, 2016). As such, the term "increased integration" refers to the erroneous binding of visual and auditory inputs that do not occur at the same time, or that is irrelevant to the the task at hand.

Age-related changes in attentional control during audiovisual perception may be an underlying mechanism behind the increased multisensory integration experienced by older adults. Attentional mechanisms facilitate the processing of reliable, task-relevant sensory inputs and inhibit/filter the processing of task-irrelevant stimuli (Pepper et al., 2023; Pepper & Nuttall, 2023; Mozolic et al., 2008; Posner & Driver, 1992; Talsma et al., 2007). Older adults find it more difficult than younger adults to initiate top-down processes against irrelevant information and hence inhibit task-irrelevant information (Zhuravleva et al., 2014; Gazzaley et al., 2005), such as ignoring background noise when trying to focus on target speech, termed the *’inhibitory deficit hypothesis’* (Hasher & Zacks, 1988; Alain & Woods, 1999). For example, after implementing an audiovisual task in which irrelevant auditory inputs had to be inhibited (cued-spatial-attention stream-bounce task), Pepper et al. concluded that older adults found it more difficult than younger adults to segregate and inhibit the task-irrelevant auditory information from being integrated with the task-relevant visual information. Older adults displayed weaker attentional control during audiovisual integration, resulting in a less accurate multisensory performance compared to younger adults.

One possible candidate mechanism for the age-related changes in attentional control during audiovisual integration may be the deployment of neural alpha oscillations. Despite historically being referred to as an "idling" rhythm associated with resting brain areas (Pfurtscheller et al., 1996; Lange et al., 2015), oscillatory alpha activity is now considered to index top-down attention (Bednar & Lalor, 2018; Wostmann et al., 2017; Sauseng et al., 2005; Capotosto et al., 2012; Thut et al., 2006) and active inhibitory processes during sensory processing (Klimesch et al., 2012; Foxe et al., 1998). Crucially, increases in alpha power over parieto-occipital areas are associated with inhibition of sensory information, preventing it from being integrated into the percept (Keller et al., 2017; Keil & Senkowski, 2017). For example, O’Sullivan et al. (2019) found that during audiovisual speech perception, when the visual information was incongruent and had to be inhibited, alpha power increased in parieto-occipital brain regions. Increases in alpha power suppressed the processing of distracting sensory inputs, to prevent the integration of incongruent auditory and visual information (Kelly et al., 2006; O’Sullivan et al., 2019). At this point, it is important to note that much of the research into the functional role of oscillatory alpha activity has been conducted on younger adult participant groups; the increased difficulty that older adults have in ignoring distracting, irrelevant sensory information (Zhuravleva et al., 2014; Gazzaley et al., 2005) may be reflected in reduced alpha power compared to younger adults during a multisensory task in which irrelevant sensory information must be inhibited.

Understanding more about the age-related changes in the attentional modulation of multisensory integration is key given our increasingly ageing population. Specifically, erroneous multisensory integration is associated with increased risk of falls in older adults (Setti et al., 2011; Stapleton et al., 2014; Mahoney et al., 2014; Peterka, 2002), as binding together task-irrelevant or incongruent sensory inputs can result in increased distractibility and inaccurate processing of relevant endogenous/exogenous stimuli (Poliakoff et al., 2006; Setti et al., 2011). Indeed, a key indicator of fall risk is the functional ability level of older adults; functional ability is often measured using composite assessments of balance ability, leg strength and gait speed. Strong functional ability is crucial for independence with healthy ageing, allowing older adults to move around the house, walk across the road, climb the stairs and perform other activities of daily living without being at a significant risk of falls (Dewhurst & Bampouras, 2014). Not only is functional ability challenged by age-related musculoskeletal decline, but can also be significantly impacted by the weaker inhibitory control of older adults. Crucially, due to their weaker attentional filtering, task-irrelevant sensory information is incorporated into older adults’ representations of their environment, which could provoke distractibility and lead to a fall (Setti et al., 2011). In other words, attention fails to reconcile the competition between relevant and irrelevant information as effectively as a function of increased age. It follows that if older adults are at an increased risk of falls (i.e. weaker balance maintenance) compared to younger adults, this may be reflected in age-related differences in alpha activity during the attentional modulation of multisensory integration, in which distracting sensory information must be suppressed.

The aims of this study were to 1) investigate the role of parieto-occipital alpha power in age-related changes in audiovisual integration, and 2) investigate the association between audiovisual integration and functional ability. Younger and older participants completed the cued-spatial-attention version of the stream-bounce task as described in Pepper et al. (2023), whilst their alpha power was extracted from parieto-occipital regions. Participants’ static and dynamic balance were also assessed. We tested the following hypotheses:

1) older adults will exhibit increased audiovisual integration compared to younger adults.

2) older adults will demonstrate weaker attentional control during audiovisual integration compared to younger adults.

3) older adults will demonstrate smaller increases from baseline in alpha power compared to younger adults.

4) balance ability will predict increased audiovisual integration and weaker attentional control during audiovisual integration

This experiment was pre-registered prior to data collection on Open Science Framework: https://doi.org/10.17605/OSF.IO/J3VPF

## Methods

### 2.1. Participants

This study included a total of 72 participants; 36 younger adults (20 males, 16 females) between 18-35 years old (*M* = 22.67, *SD* = 4.09) and 36 older adults (14 males, 22 females) between 60-80 years old (*M* = 66.86, *SD* = 4.43). This sample size was determined via an a-priori power analysis using the pwr package in R studio (see pre-registration on Open Science Framework: https://doi.org/10.17605/OSF.IO/J3VPF). Specifically, the pwr.f2.test function was implemented as recommended for multiple regression/general linear model analyses (Kabacoff, 2015), using the large effect size generated by Pepper et al. (2023) and Kelly et al. (2006), a numerator degrees of freedom of 14, an alpha significance level of 0.05 and a power of 80%.

Participants were eligible for the study if they considered themselves fluent English speakers with normal or corrected-to-normal vision, screened for via self-report. Participants were ineligible to participate if they had a history or current diagnosis of cognitive impairments or neurological conditions (e.g. epilepsy, mild cognitive impairment, dementia, Parkinson’s Disease) or learning impairments (e.g. dyslexia). Participants were also ineligible to participate if they had moderate-severe hearing loss resulting in the wearing of hearing aids; if they suffered from motion sickness; if they were diagnosed with any vestibular impairments (e.g. vertigo) or numbness in the lower limbs; if they were diagnosed with any muscle or bone conditions which could prevent standing comfortably (including lower limb, hip or spine surgery within the last year, or recent injury); if they relied on assistive walking devices (e.g. canes or walking frames), or if they were on medication which depresses the nervous system or affects balance (Thomas et al., 2016).

Participants were recruited via opportunity sampling; younger participants were students at Lancaster University, whilst older participants were recruited through the Centre for Ageing Research at Lancaster University; through advertising to local community groups, such as University of the Third Age; or through word of mouth. All participants provided informed consent. Ethical approval was received from Lancaster University Faculty of Science and Technology Ethics Committee (ref: FST-2022-0636-RECR-3).

### 2.2. Pre-screening tools

Participants were asked to complete two pre-screening questionnaires using Qualtrics online platform (Qualtrics XM, Provo, UT), to assess their eligibility for the study prior to coming to the lab.

#### 2.2.1. Speech, Spatial and Quality of Hearing Questionnaire (SSQ; Appendix A; Gatehouse & Noble, 2004)

Participants rated their hearing ability in different acoustic scenarios using a sliding scale from 0-10 (0=“Not at all”, 10=“Perfectly”). Whilst, at present, no defined cut-off score on the SSQ is available as a parameter to inform decision-making, previous studies have indicated that a mean score of less than 5.5 is indicative of moderate hearing loss (Gatehouse & Noble, 2004). As a result, people whose average score on the SSQ was lower than 5.5 were not eligible to participate in the experiment. This was to ensure that any changes in audiovisual integration measured in the task would not be due to a participant’s inability to hear the auditory stimuli. Hearing acuity was then evaluated objectively using pure-tone audiometry (see section 2.2.3) when participants attended the lab.

#### 2.2.2. Informant Questionnaire on Cognitive Decline in the Elderly (IQ-CODE; Appendix B; Jorm, 2004)

Participants used a self-report version of the IQ-CODE to rate how their performance in certain tasks has changed compared to 10 years ago, answering on a 5-point Likert scale (1=“Much Improved”, 5=“Much worse”). An average score of 3.65 is the usual cut-off point when evaluating cognitive impairment and dementia (Slade et al., 2023; Jansen et al., 2008), therefore people whose average score was higher than 3.65 were not eligible to participate in the experiment. This was to ensure that any changes in audiovisual integration measured in the task would not be due to the participant experiencing mild cognitive impairment.

#### 2.2.3 Pure-Tone Audiometry

If the online SSQ and IQCODE pre-screening questionnaires deemed the participants eligible for the study, they were invited to the lab for the in-person testing session. Pure-tone thresholds were measured bilaterally at 0.25 kHz, 0.5 kHz, 1 kHz, 2 kHz, 4 kHz and 8 kHz, in accordance with the British Society of Audiology (2018) guidelines. Pure tone average thresholds were averaged across 0.5-4kHz in each ear, and then averaged across ears. Audiometry was used to ensure that any differences in multisensory performance were not due to moderate-severe hearing loss. The mean pure-tone audiometry thresholds for each age group are displayed in *Figure 1*.

**Figure 1.**
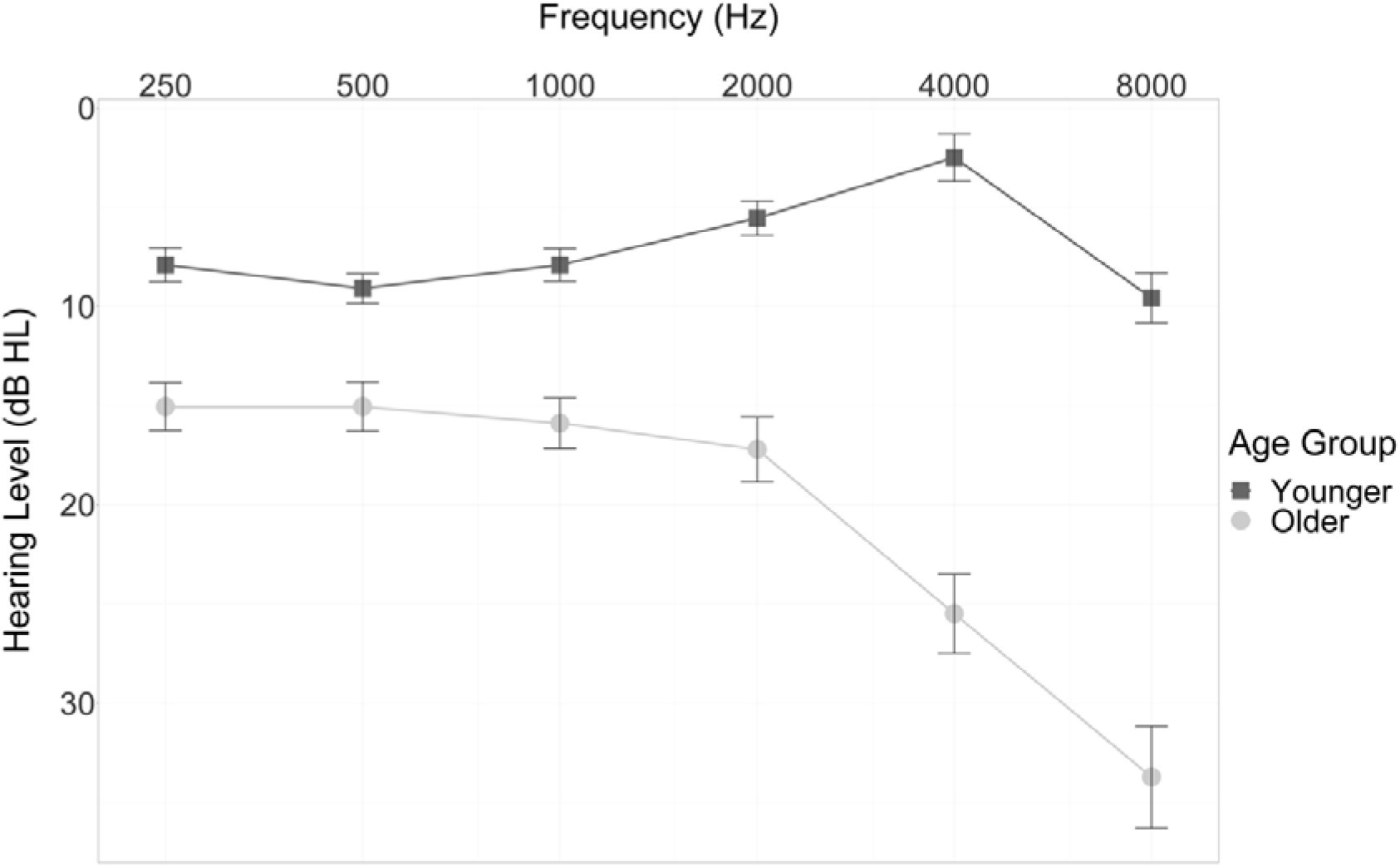
Mean pure-tone audiometry thresholds recorded for each age group at each frequency. Black markers represent data of younger adults, grey markers represent the data of older adults. Standard error displayed as error bars.

The mean scores of eligible participants in each pre-screening assessment are summarised in *Table 1*. Independent *t*-tests revealed there was no significant difference between age groups on the SSQ [*t*(70) = -0.92, *p*=.154; *M_Younger_* = 8.43, *M_Older_* = 8.64]. Older adults scored significantly higher score on the IQ-CODE questionnaire compared to younger adults [*t*(70) = -11.50, *p*<.001; *M_Younger_* = 1.96, *M_Older_* = 3.07]. Older adults had significantly higher PTA thresholds compared to younger adults [*t*(70) = -8.16, *p*<.001, *M_Younger_* = 6.27, *M_Older_* = 18.30].

**Table 1.** Mean scores on the *Speech, Spatial and Quality of Hearing Questionnaire* (SSQ), *Informant Questionnaire on Cognitive Decline in the Elderly* (IQ-CODE) and pure-tone audiometry (PTA) pre-screening measures, for both younger and older adults. Data is presented as mean (SD). Significance was set at a p<0.05.

| **Test** | **Younger** | **Older** | ***p*-value** |
| --- | --- | --- | --- |
| **SSQ** | 8.43<br>(0.91) | 8.64<br>(1.09) | $p = .154$ |
| **IQCODE** | 1.96<br>(0.55) | 3.07<br>(0.19) | $p < .001$ |
| **PTA** | 6.27<br>(4.56) | 18.30<br>(7.59) | $p < .001$ |

### 2.3. Experimental Design

#### 2.3.1. Questionnaire Measures

After passing the pre-screening eligibility assessments, all participants completed two self-report assessments of physical activity, providing detailed information regarding participants’ own perception of their balance abilities and their fitness levels.

##### 2.3.1.1 Activities-Based Balance Confidence Scale (ABC; Appendix C; Powell & Myers, 1995)

The ABC scale is a 16-item questionnaire used to assess participants’ balance confidence in performing daily activities. Participants were asked to rate how confident they are in performing each activity, on a 10-point scale ranging from 0% (not confident at all) to 100% (completely confident). An average score of greater than 80% indicates high levels of functioning; a score of between 50% and 80% indicates moderate levels of functioning; a score of less than 50% indicate low levels of functioning. Crucially, a score of less than 67% is indicative of a substantial risk of falling.

##### 2.3.1.2. Rapid Assessment of Physical Activity (RAPA; Appendix D; Topolski et al., 2006)

RAPA is a 9-item questionnaire used to assess the level of physical activity in our participants. Participants are asked to answer Yes/No to whether the physical activity level in the scenario accurately describes them. The scale is divided into two parts. RAPA1 consists of 7 items and measures cardio-respiratory, aerobic activity (scored as 1 = "Sedentary", 2 = "underactive", 3 = "underactive regular - light activities", 4 and 5 = "underactive regular", 6 and 7 = "Active"). The highest affirmative score provided by participants is their final recorded score for RAPA1. RAPA2 consists of 2 items and measures strength and flexibility-based physical activity. An affirmative response to the first item results in a score of 1; an affirmative response to the second item results in a score of 2; affirmative responses to both items scores 3; negative responses to both items scores 0. Participants’ scores on RAPA1 and RAPA2 were added together to provide an overall indication of physical activity levels of the samples. Higher total scores represent higher levels of physical activity.

#### 2.3.2 Functional Ability – The Short Physical Performance Battery (SPPB; Guralnik et al., 1994, 2000)

The SPPB is divided into three sections measuring balance, gait speed and leg strength, each of which are scored from 0-4 and added together to provide a composite measure of functional ability. As a result, the minimum score on the SPPB was 0 points and the maximum score was 12 points. Lower scores on the SPPB are indicative of weaker lower-body functioning and an increased risk of falls (Guralnik et al., 2000).

To increase the sensitivity of the data collected in these physical assessments, force platforms were implemented during the standing balance stage of the SPPB. Participants were asked to stand on force platforms with feet side-by-side, in a semi-tandem position, and in a tandem position, for 10 seconds each (if able to). Force platforms (PASCO, Roseville, CA, USA) collected centre of pressure movements in the anteroposterior and mediolateral axis, which were used to calculate sway area and sway velocity in each of the stance conditions. The force platforms were positioned side by side, without touching each other and recorded at a rate of 100Hz. Participants were asked to keep their hands by their sides throughout each assessment and focus on the wall ahead of them. Sway area and sway velocity values from the three stances were averaged and used for further analysis. SPPB scores were therefore considered to be measures of overall functional ability, whilst the sway measures extracted during the SPPB were considered to be measures of balance specifically.

#### 2.3.3 Timed Up and Go Test (TUG; Podsiadlo & Richardson, 1991; Shumway-Cook et al., 2000)

As an additional dynamic measure of functional ability, participants were also asked to complete the Timed Up-And-Go (TUG) test, which is a clinical assessment of fall risk in older adults. Participants are asked to stand from the chair, walk 3 metres at a comfortable pace, turn around, walk back to the chair and sit down. The time that participants took to complete this assessment was recorded, with longer times (greater than 13.5 seconds for community-dwelling older adults; Shumway-Cook et al., 2000) indicating increased fall risk.

#### 2.3.4 The Stream-Bounce Task

This behavioural task implemented a 2 (Age: Younger vs Older) x 2 (Cue: Valid vs Invalid) x 3 (Stimulus Onset Asynchrony [SOA]: Visual Only [VO] vs 0 milliseconds vs 300 milliseconds) mixed design, with Age as a between-subjects factor and Cue and SOA as within-subjects factors.

The stream-bounce stimuli used in the task were replicated from Donohue et al. (2015), with experimental details described previously in Pepper et al. (2023). Briefly, at the start of each trial, participants were cued either towards the full "X" shaped motion of the stimuli (validly-cued trials) or towards the stopped motion of the stimuli (invalidly-cued trials) appearing on the computer screen. Two thirds of the trials contained a task-irrelevant sound, played either synchronously with the circles intersecting (0ms delay) or 300ms afterwards. The remaining trials were visual-only. At the end of each trial, participants were asked whether they perceived the circles to "pass through" or "bounce off" each other.

The experiment consisted of 12 different trial conditions, randomised across all participants. The experimental block contained of a set of 60 validly-cued trials and a set of 60 invalidly-cued trials (two conditions), which were equally distributed between each side of the screen (left/right) and three stimulus onset asynchrony (SOA) conditions (Visual Only [VO], 0 milliseconds and 300 milliseconds); this means that each participant completed 120 valid trials and 120 invalid trials for each SOA. Participants completed the experiment in a quiet room on an Apple Mac computer (version 12.2.1) with a standard keyboard. All participants wore EEG-compatible earphones (ER2 ultra-shielded insert earphones; Intelligent Hearing Systems). A volume check was conducted at the beginning of the experiment; participants were presented with a constant tone and the volume of this tone was adjusted to a clear and comfortable level. Screen captures of a validly-cued, 0ms SOA trial are displayed in *Figure 2.* The percentage of “Bounce” responses provided in each Cue x SOA condition was calculated for each participant.

**Figure 2.**
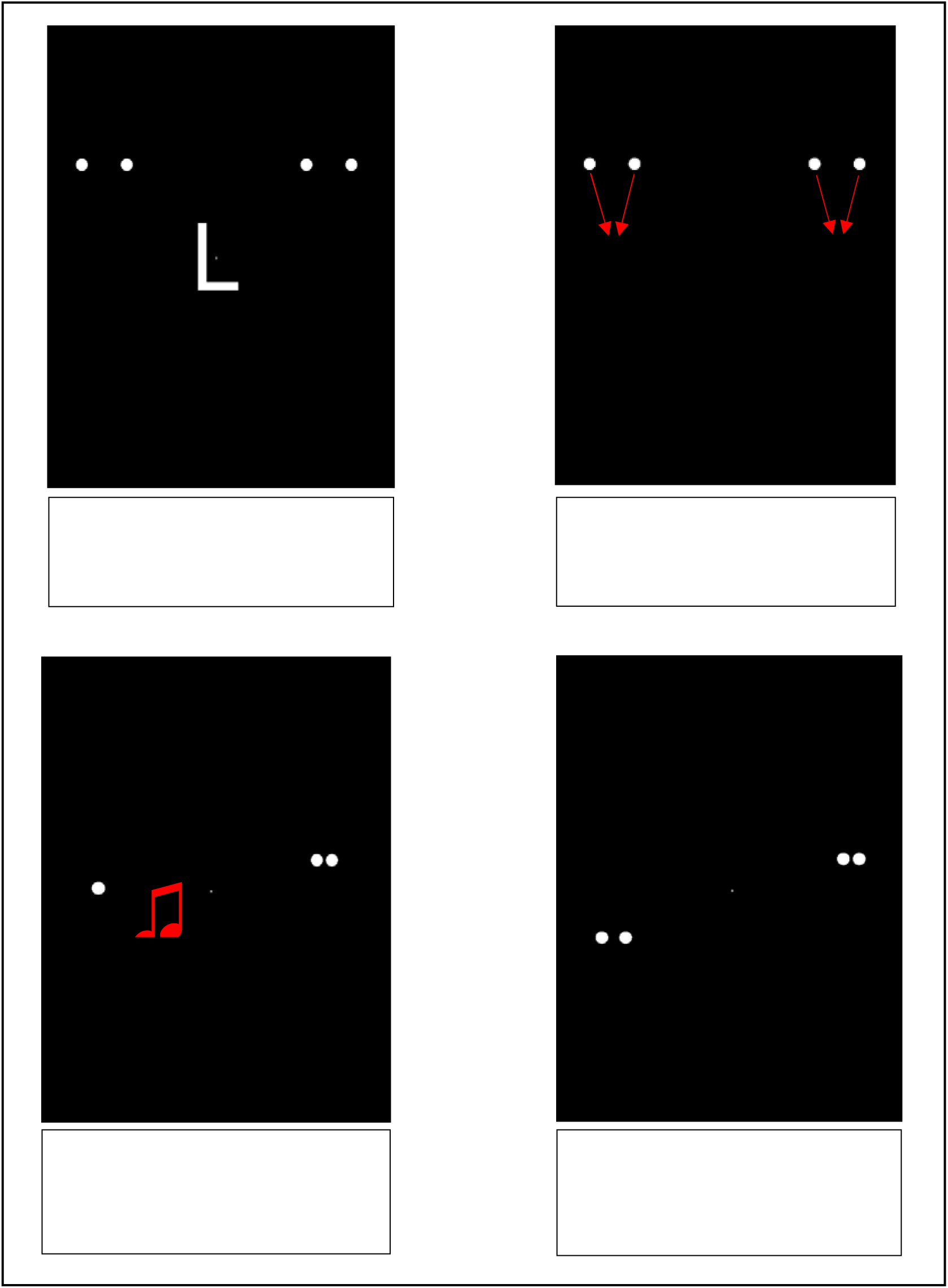
Screen captures of a validly-cued trial (valid left), with an SOA of 0ms (sound synchronous with intersection). Participants provided their pass/bounce judgement at the end of the trial. Image taken from the published manuscript of Pepper et al. (2023).

#### 2.3.5 EEG Data Acquisition and Pre-Processing

Continuous EEG data were sampled at 500Hz from a 32-channel EEG amplifier system (BrainAmps, BrainProducts GmbH, Germany) with Ag/AgCl electrodes positioned according to the international 10-20 system (actiCAP EasyCap, BrainProducts, GmbH, Germany), referenced to the central Reference electrode during recording. The data underwent online bandpass filtering, applying a low cut-off filter of 0.1Hz, a high-cut-off filter of 40Hz, and a notch filter of 50Hz. Psychopy and BrainVision Recorder (version 1.10, Brain Products GmbH, Germany) were used in conjunction to record trial-specific information in real time, including EEG triggers coded to identify the condition each participant experienced and when the participant provided a key press response (Franzen et al., 2020; Klatt et al., 2020). These data were collected and stored for offline analysis in EEGLAB.

Processing and EEG analyses were completed offline using the EEGLAB toolbox (Delorme & Makeig, 2004) and MATLAB scripts. The EEG data was first resampled to 256Hz and re-referenced to the average of all electrodes. Breaks between experimental blocks were removed and an independent component analysis (ICA) was performed on the data. Artefactual independent components were detected and rejected using the ICFlag function in EEGLAB; components that were identified as being over 80% likely to be heart, muscle or eye artefacts were removed from the dataset (Delorme et al., 2007). The pre-processed EEG data was then epoched, beginning at the presentation of the fixation cross in the stream-bounce task and ending 3 seconds afterwards once the circles had completed their full motion.

#### 2.3.6 Alpha power extraction

Alpha power was extracted from the 8-12Hz frequency band at electrodes positioned over the parietal and occipital lobes (P3, P4, P7, P8, O1, O2, Oz). The use of parieto-occipital electrodes is in line with previous research investigating posterior alpha activity for audiovisual integration (Getzmann et al., 2020; O’Sullivan et al., 2019; van Driel et al., 2017; Thut et al., 2006; Klatt et al., 2020). Alpha power was determined using the power spectral density (PSD) package in EEGLAB. The ’spectopo’ function is based upon Welsch’s method and uses a 256-point Hamming window. Within each epoch, for each participant, mean alpha power over each electrode was calculated for the 1000ms pre-stimulus interval of each condition type, and for the 2000ms stream-bounce trial of each condition type. The alpha power was then averaged across all electrodes of interest, to produce a grand mean alpha power value for the experimental condition, and a grand mean alpha power for the pre-stimulus baseline associated with each condition. Mean experimental alpha power was then subtracted from mean baseline alpha power to produce an alpha power value representative of the difference in alpha power between ‘rest’ (pre-stimulus interval) and the experimental trial.

The procedure outlining the entirety of the study is displayed in *Figure 3*.

**Figure 3.**
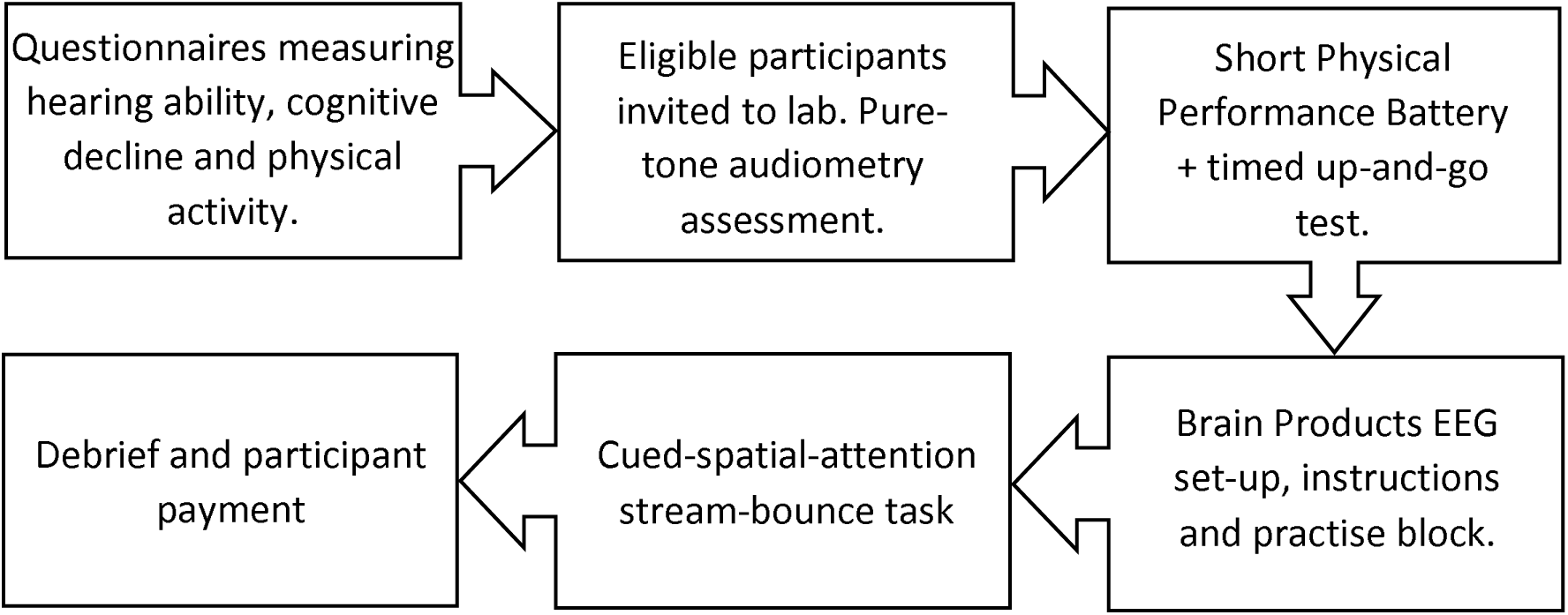
Flowchart detailing the procedure of the study.

### 2.4. Statistical Analyses

Two multiple linear regression models were run to examine whether a) age, oscillatory alpha power and balance ability and the interactions of each variable with age can predict audiovisual integration in the stream-bounce task (Model 1: Proportion of "Bounce" responses = Age + Cue + SOA + Alpha power difference + Sway Velocity + SPPB Score + Pure-tone audiometry + Age*Cue + Age*Alpha + Age*Velocity + Age*SPPB Score), and b) age, audiovisual integration and balance ability and the interactions of each variable with age can predict oscillatory alpha power (Model 2: Alpha power difference = Age + Cue + SOA + Bounce + Sway Velocity + SPPB Score + Pure-tone audiometry + Age*Cue + Age*Bounce + Age*Velocity + Age*SPPB Score). Prior to the examination of the models, the variables were assessed for violation of the related assumptions; the assessment confirmed that all relevant assumptions were met. To correct for multiple models, all regression analyses were conducted using an alpha value of p=.025 and the adjusted values are reported. To address the violation of ANOVA assumptions present with percentage data, these grand means were converted into z-scores, following the procedures recommended by Caldwell et al. (2019).

Data were analysed and visualised in R Studio (version 4.2.1) using the ’stats’ (R Core Team, 2022), ’car’ (Fox & Weisberg, 2019), ’performance’ (Ludecke et al., 2021), ’emmeans’ (Lenth, 2023) and ’ggplot2’ (Wickham, 2016) packages. Post-hoc ANOVAs and correlational analyses were used to analyse the differences and relationships between conditions and age groups. Pure-tone audiometry thresholds were included within each model to control for any age-related differences in hearing ability.

### 2.5. Deviations from pre-registration

In the pre-registration for this study, it was proposed to include the TUG test times in both models as a measure of functional ability. However, as is indicated by the very high ABC and RAPA scores (see *Table 4*), the older adult sample in this study were very physically able. As a result, the TUG test is not likely to be sensitive enough to detect fall risk in these active older adults (Barry et al., 2014), while not allowing separation of the different elements that contribute to its performance. Given that the SPPB can also be used as a measure of functional ability, while the distinct and distinguishable measures it comprises of allows direct assessment of balance, it was deemed unnecessary to include both the SPPB and the TUG in the model. As such, and after finding moderate collinearity between Sway Area and Sway Velocity during model checks, Sway Area and Timed Up-And-Go times were omitted as model predictors.

## Results

### H1 – Older adults will exhibit increased audiovisual integration compared to younger adults

The mean proportion of “Bounce” responses within each condition, for each age group, are displayed in *Figure 4*.

**Figure 4.**
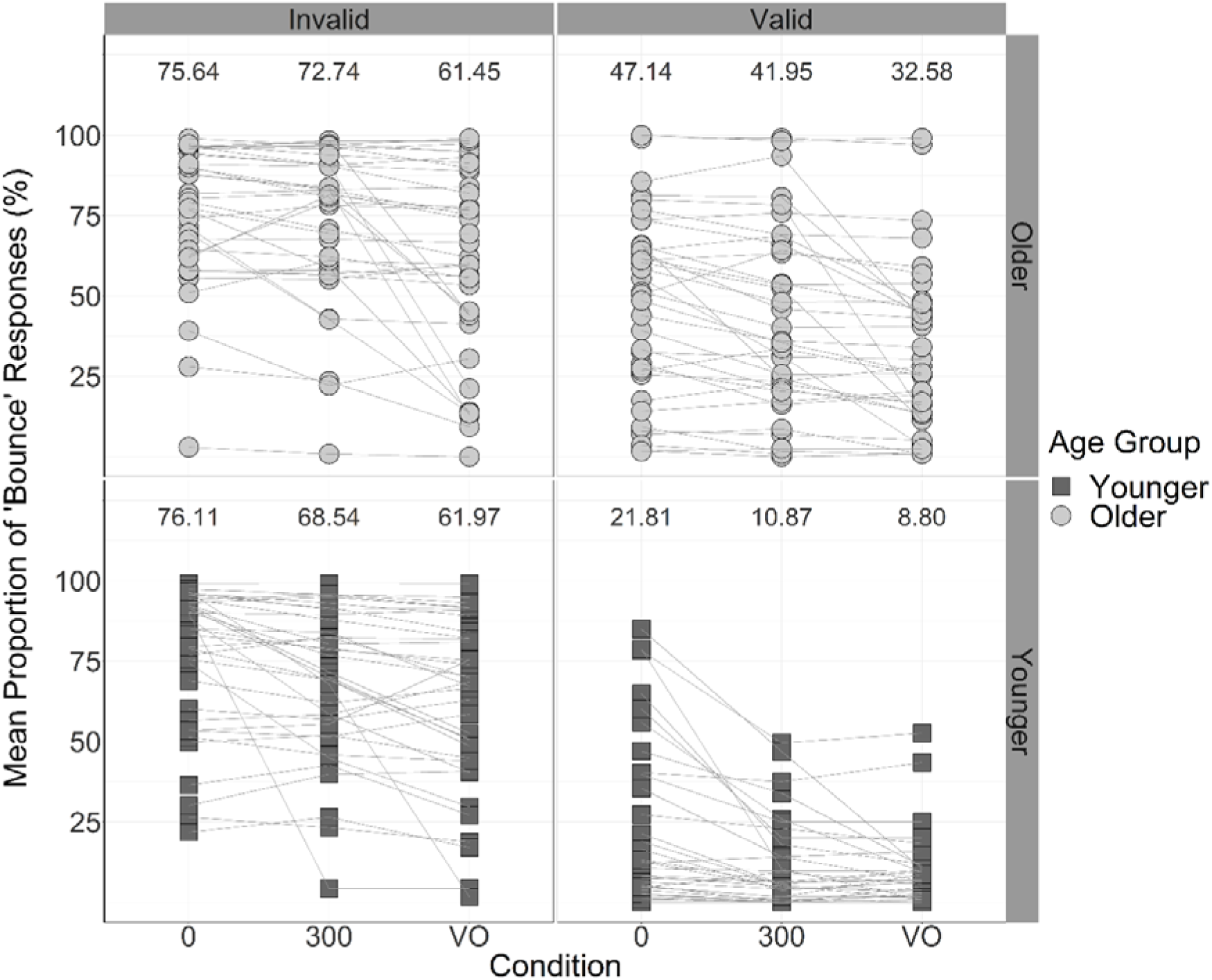
Mean proportion of “Bounce” responses in each Cue and SOA condition for each participant. Bottom panels represent data of younger adults, top panels represent the data of older adults. Participants’ “Bounce” responses are linked across conditions using lines. Numbers at the top of each panel display the mean proportion of “Bounce” responses in each condition.

For Model 1, the outcome was the proportion of "Bounce" responses produced in the stream-bounce task. The model was significant overall [F(12,413) = 36.47, p<.001, adjusted R^2^ = 0.50]. The output from the ANOVA performed on the multiple regression model is displayed in Table 2.

**Table 2.** Multiple linear regression output detailing the statistical contribution of each prediction and interaction to the outcome of the proportion of "Bounce" responses.

| Full Model: Bounce = Age + Cue + SOA + Alpha + Velocity + SPPB Score + PTA + Age*Cue + Age*Alpha + Age*Velocity + Age*SPPB Score |  |  |  |  |  |  |
| --- | --- | --- | --- | --- | --- | --- |
| Predictors | Df | Sum of squares | Mean square | *F value* | *p* | *Adjusted p-value* |
| **Age Group** | **1** | **16.64** | **16.64** | **33.33** | **<.001** | **<.001** |
| **Cue** | **1** | **167.26** | **167.26** | **335.01** | **<.001** | **<.001** |
| **SOA** | **2** | **12.31** | **6.16** | **12.33** | **<.001** | **<.001** |
| Alpha | 1 | 0.06 | 0.06 | 0.11 | 0.741 | 1.00 |
| Sway Velocity | 1 | 1.97 | 1.97 | 3.95 | 0.048 | 0.095 |
| SPPB Score | 1 | 2.48 | 2.48 | 4.96 | 0.026 | 0.053 |
| PTA | 1 | 0.15 | 0.15 | 0.29 | 0.589 | 1.00 |
| **Age\*Cue** | **1** | **15.79** | **15.79** | **31.63** | **<.001** | **<.001** |
| Age*Alpha | 1 | 0.02 | 0.02 | 0.05 | 0.830 | 1.00 |
| Age*Velocity | 1 | 1.29 | 1.29 | 2.59 | 0.108 | 0.217 |
| Age*SPPB Score | 1 | 0.55 | 0.55 | 1.09 | 0.296 | 0.592 |
| Residuals | 413 | 206.20 | 0.50 |  |  |  |

*Notes: Df = degrees of freedom; SOA = Stimulus-Onset Asynchrony; SPPB = Short Physical Performance Battery; PTA = pure-tone audiometry*

With regards to the individual predictors in the model, there was a significant main effect of Age on the proportion of "bounce" responses [*F*(1, 413) = 33.33, *p*<.001]. Overall, a significantly greater proportion of "bounce" responses were produced by older adults (*M* = 55.25%, *SE* = 2.21, 95% CI [50.85, 59.65]) than by younger adults (*M* = 41.35%, *SE* = 2.21, 95% CI [36.95, 45.75]; mean difference = 13.90%, *SE* = 3.12, 95% CI [7.68, 20.12]), providing support for hypothesis one which predicted that older adults will exhibit increased audiovisual integration compared to younger adults.

### H2: Older adults will demonstrate weaker attentional control during audiovisual integration compared to younger adults

The interaction between Age and Cue was a significant predictor of "Bounce" responses [*F*(1, 419) = 31.41, *p*<.001]. To analyse pairwise comparisons within the Age and Cue interaction, the “bounce” responses in each SOA condition were collapsed, so that a mean percentage of “bounce” responses provided by each participant could be calculated for validly cued and invalidly cued conditions. These percentages were then converted to standardized z-scores.

#### Age pairwise comparisons

To assess differences between the proportion of “bounce” responses provided by younger adults and older adults in valid trials, and the differences between younger and older adults in invalid trials, two separate one-way ANOVAs were conducted. The first one-way ANOVA analysed responses in the valid condition, and revealed that there were significant differences in the proportion of “bounce” responses between age groups, [*F*(1, 70) = 27.61, *p* < .001]. In the valid condition, a significantly greater proportion of “bounce” responses were produced by older adults (*M* = 40.56%, *SE* = 4.39, 95% CI [31.64, 49.47]) than younger adults (*M* = 13.83%, *SE* = 2.57, 95% CI [8.61, 19.04]). This provides support for hypothesis two, suggesting that even in the validly-cued conditions, older adults still integrated the visual and auditory information more frequently than younger adults.

The second one-way ANOVA analysed responses in the invalid condition, and in contrast, indicated no significant difference between age groups, [*F*(1, 70) = 0.04, *p* = .839]. In the invalid condition, a similar proportion of “bounce” responses were produced by younger adults (*M* = 68.87%, SE = 3.62, 95% CI [61.52, 76.23]) and by older adults (*M* = 69.94%, *SE* = 3.78, 95% CI [62.27, 77.61]).

#### Cue pairwise comparisons

To assess differences in the proportion of “bounce” responses provided by younger adults in valid versus invalid trials, and by older adults in valid versus invalid trials, a repeated-measures ANOVA was conducted on the collapsed z-score data. When examining the data of younger adults, there was a significant difference in the proportion of “bounce” responses in validly cued and invalidly cued trials, [*F*(1, 35) = 155.44, *p* < .001, η_p_ ^2^ = 0.82]. Overall, younger adults produced a significantly greater proportion of “bounce” responses in invalidly cued trials (*M* = 68.87%, *SE* = 3.62, 95% CI [61,52, 76.23]) compared with validly cued trials (*M* = 13.83%, *SE* = 2.57, 95% CI [8.61, 19.04]; mean difference = 55.05%, *SE* = 4.42). When examining the data of older adults, there was also a significant difference in the proportion of “bounce” responses in the validly cued and invalidly cued trials, [*F*(1, 35) = 17.93, *p* < .001, η_p_ ^2^ = 0.34]. Overall, older adults produced a greater proportion of “bounce” responses in the invalid trials (*M* = 69.94%, *SE* = 3.78, 95% CI [62.27, 77.61]) compared to valid trials (*M* = 40.56%, *SE* = 4.39, 95% CI [31.64, 49.47]; mean difference = 29.39%, *SE* = 6.94). Taken together, these results confirmed hypothesis 2 in that older adults exhibited weaker attentional filtering during audiovisual integration compared to younger adults.

### H3: Older adults will show smaller increases from baseline in alpha power compared to younger adults

For Model 2, the outcome was the difference in alpha power from baseline in the experimental trials of the stream-bounce task. The model was significant overall [*F*(12,413) = 2.03, *p*=.021, adjusted R^2^ = 0.03]. There was no significant main effects in the model, and the interactions between age and cue, age and "bounce" responses and age and velocity were not significant. As a result, the data did not support hypothesis three that alpha power would reflect age-related changes in attentional control during multisensory integration. However, there was a significant interaction between age and SPPB scores on alpha power [*F*(1, 413) = 17.86, *p*<.001]. The output of the ANOVA conducted on the multiple regression model is displayed in Table 3.

**Table 3.** Multiple linear regression output detailing the statistical contribution of each prediction and interaction to the outcome of alpha power.

| Full Model: Alpha ~ Age + Cue + SOA + Bounce + Velocity + SPPB Score + Age*Cue + Age*Bounce + Age*Velocity + Age*SPPB Score |  |  |  |  |  |  |
| --- | --- | --- | --- | --- | --- | --- |
| Predictors | Df | Sum of squares | Mean square | F value | p | Adjusted p-value |
| Age Group | 1 | 4.26 | 4.26 | 1.89 | 0.170 | 0.340 |
| Cue | 1 | 0.53 | 0.53 | 0.24 | 0.628 | 1.00 |
| SOA | 2 | 0.38 | 0.19 | 0.08 | 0.920 | 1.00 |
| Bounce | 1 | 0.23 | 0.23 | 0.10 | 0.747 | 1.00 |
| Sway Velocity | 1 | 0.37 | 0.37 | 0.16 | 0.686 | 1.00 |
| SPPB Score | 1 | 1.76 | 1.76 | 0.78 | 0.377 | 0.755 |
| PTA | 1 | 0.01 | 0.01 | 0.01 | 0.936 | 1.00 |
| Age* Cue | 1 | 0.05 | 0.05 | 0.02 | 0.882 | 1.00 |
| Age*Bounce | 1 | 0.70 | 0.70 | 0.31 | 0.578 | 1.00 |
| Age*Velocity | 1 | 7.51 | 7.51 | 3.33 | 0.069 | 0.756 |
| **Age\*SPPB Score** | **1** | **39.00** | **39.00** | **17.29** | **<.001** | **<.001** |
| Residuals | 413 | 931.25 | 2.26 |  |  |  |

*Notes: Df = degrees of freedom; SOA = Stimulus-Onset Asynchrony; SPPB = Short Physical Performance Battery score; PTA = pure-tone audiometry threshold*

### H4: Balance will predict audiovisual integration and attentional control

After correcting p-values for multiple comparisons, there was no significant main effect of SPPB score on the proportion of "Bounce" responses [Model 1, Table 2; *F*(1, 413) = 4.96, *p*=0.291]. In addition, after correcting p-values, there was no significant main effect of sway velocity on the proportion of "bounce" responses [*F*(1, 413) = 3.95, *p*=.524]. Taken together, the data did not support hypothesis four that weaker functional ability or balance ability would predict audiovisual integration within the stream-bounce task.

However, Model 2 (Table 3), indicated there was a significant interaction between age and SPPB scores on alpha power [*F*(1, 413) = 17.86, *p*<.001]. To analyse this interaction, correlational analyses were conducted, assessing the relationship between alpha power and SPPB scores in younger adults and in older adults. These exploratory correlational analyses revealed that for younger adults, there was a significant negative relationship between alpha power and SPPB scores [*r*(214) = -0.15, *p*=.025], with higher alpha power being associated with weaker balance ability. In contrast, for older adults, there was a significant positive relationship between alpha power and SPPB scores [*r*(214) = 0.20, *p*=.002], with higher alpha power being associated with stronger functional ability. Participants SPPB scores and sway velocities are displayed in Figures 5 and 6.

**Figure 5.**
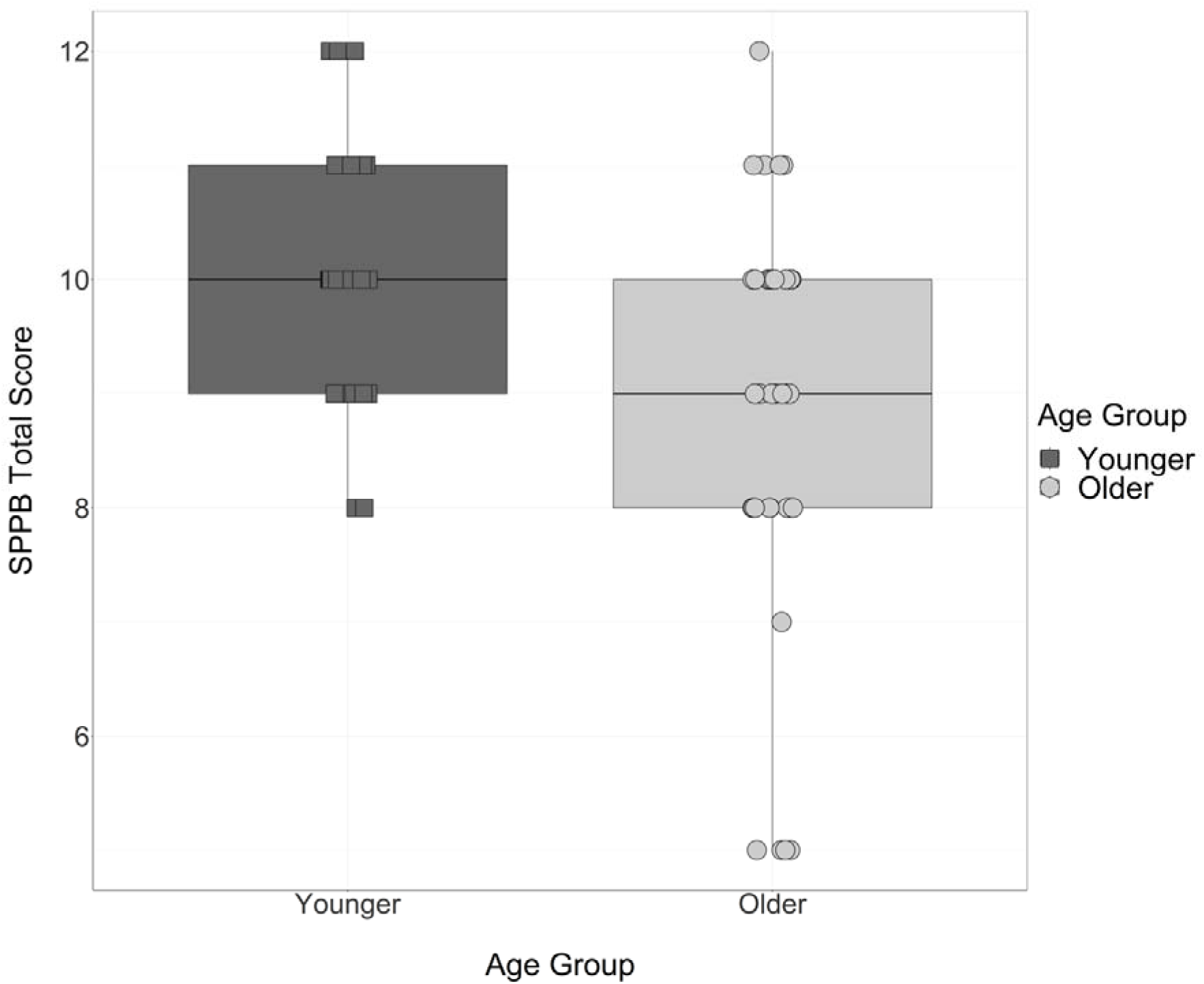
Short Physical Performance Battery scores for all participants. Black squares and boxplot represent data of younger adults, grey circles and boxplot represent the data of older adults. Each boxplot displays the median, the lower and upper quartile for each condition.

**Figure 6.**
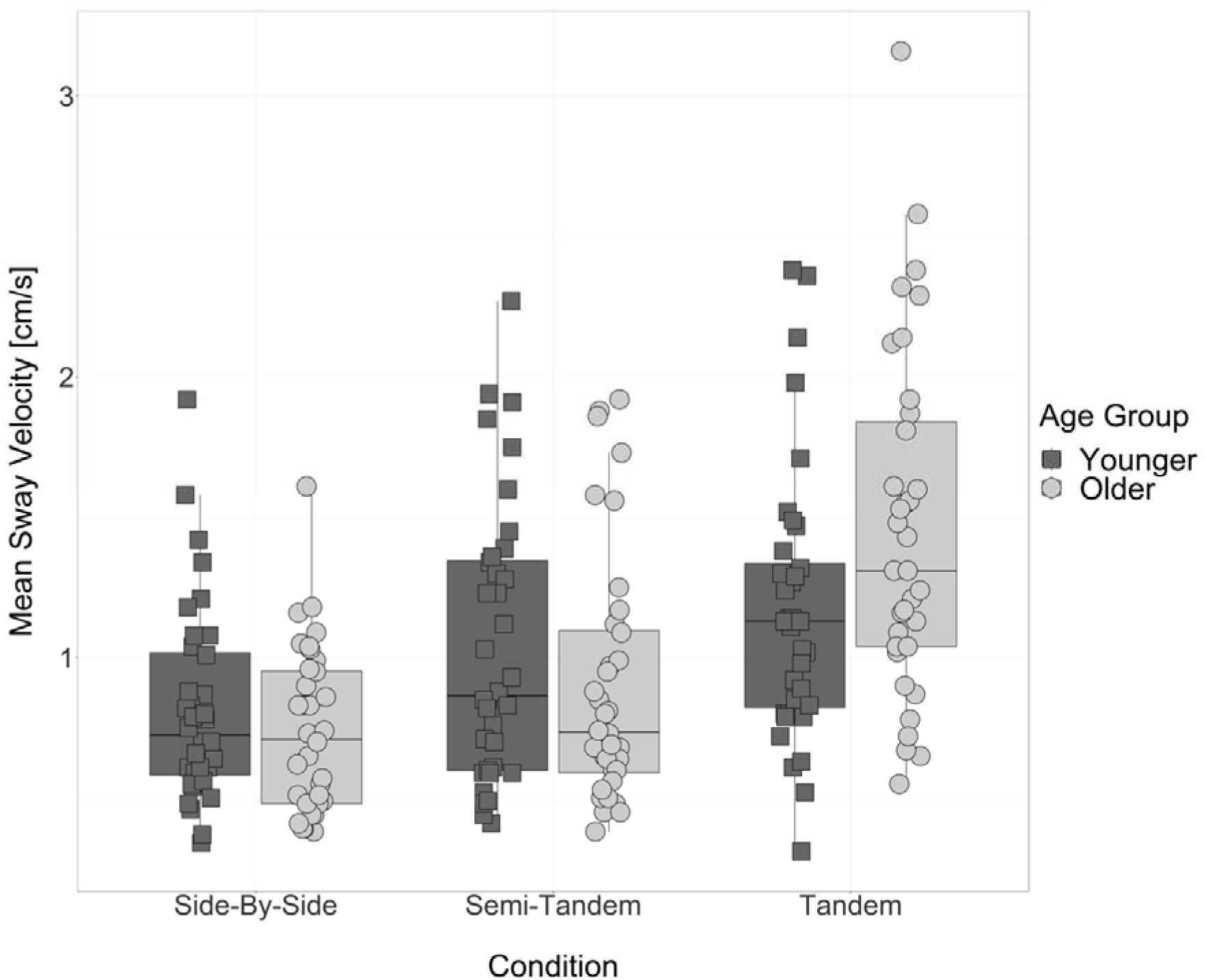
Mean sway velocity recorded for each participant in the side-by-side, semi-tandem and tandem balance positions in the Short Physical Performance Battery. Black squares and boxplots represent data of younger adults, grey circles and boxplots represent the data of older adults. Each boxplot displays the median, the lower and upper quartile for each condition.

Exploratory analyses were also conducted to investigate differences between age groups for their subjective perspectives of their balance ability and physical activity levels, using the questionnaire data collected from participants before the testing session. The mean scores on the ABC and RAPA questionnaires are displayed in *Table 4*, with individual scores displayed in *Figure 7* and *Figure 8*. An independent *t*-test revealed that there was no significant difference between age groups on the ABC [*t*(70) = 0.48, *p*=.995] or on total RAPA scores [*t*(70) = 0.63, *p*=.282].

**Table 4.** Mean scores on the ABC and RAPA self-report questionnaires on balance confidence and physical activity, for both younger and older adults. Standard deviations displayed in parentheses.

| **Age group** | **ABC** | **RAPA1**<br>**(aerobic)** | **RAPA2**<br>**(strength)** | **ABC Class** | **RAPA Class** |
| --- | --- | --- | --- | --- | --- |
| Younger | 95.10<br>(9.61) | 5.67<br>(1.59) | 1.42<br>(1.11) | High<br>functioning | Underactive<br>regular |
| Older | 94.16<br>(6.72) | 5.72<br>(1.16) | 1.06<br>(1.24) | High<br>functioning | Underactive<br>regular |

## Discussion

The aims of this study were to 1) investigate the role of parieto-occipital alpha power in age-related changes in audiovisual integration, and 2) investigate the association between audiovisual integration and functional ability. Whilst results from the stream-bounce task provide support for the theory that older adults exhibit increased audiovisual integration and weaker attentional control compared to younger adults, oscillatory alpha power, functional ability and balance ability did not predict such changes. However, an interaction between participants’ functional ability and age predicted alpha power within the task; this indicates younger and older adults may display a differential reliance on attentional mechanisms for functional ability.

### Older adults displayed weaker attentional control during audiovisual integration compared to younger adults

The finding that older adults produced a greater proportion of "Bounce" responses compared to younger adults, even when attending to the validly-cued location, is consistent with the results of Pepper et al. (2023) and supports our hypothesis. Despite the fact that participants were paying attention to the visual stimuli in the validly-cued condition, which should strengthen the ability to suppress the task-irrelevant sound, older adults integrated the sound and the visual intersection more frequently than younger adults did. This is in line with the inhibitory deficit hypothesis (Hasher & Zacks, 1988) – older adults found it more difficult to inhibit the distracting sound in the stream-bounce task and as a result, produced a greater proportion of “Bounce” responses, even if the sound occurred after the circles intersected. This kind of erroneous multisensory integration exhibited by older adults has important consequences for their ability to safely perceive and navigate through dynamic environments, in that weaker top-down modulation of multisensory integration may result in the increased processing of irrelevant sensory information. It is important that future research investigates complex audiovisual stimuli that participants encounter in real-world environments (e.g. speech). This will allow researchers to arrive at more ecologically valid conclusions regarding the impact of older adults’ increased integration and weaker attentional control on perception and action. The use of dynamic visual stimuli, however, is useful for studying the impact of age-related changes in audiovisual integration on fall risk, due to the importance of optic flow mechanisms, for example, in guiding safe locomotion and maintaining balance (Raffi & Piras, 2019; Peterka et al., 1995).

### Oscillatory alpha power did not predict audiovisual integration

The data in the current study did not provide support for our hypothesis – alpha power did not predict the proportion of "Bounce" responses produced in the stream-bounce task. A potential reason for this is that perhaps analysing alpha activity alone is insufficient for investigating the interplay between multisensory integration and inhibitory control (Talsma et al., 2010), especially when the moving stimuli used in this task are more complex than simple flashes and beeps. For example, whilst alpha activity appears to be crucial in top-down attentional control and inhibitory functioning, gamma activity (30-80Hz) is believed to reflect the bottom-up processing of low-level sensory inputs (Keil & Senkowski, 2018; Krebber et al., 2015; Scurry et al., 2021). Scurry et al. (2021) implemented the sound-induced flash illusion with younger, healthy older and fall-prone older adults, measuring their alpha and gamma activity throughout. The researchers found that fall-prone older adults were more susceptible to the sound-induced flash illusion, displaying increased integration and less accurate multisensory perception. Importantly, these fall-prone older adults displayed reduced phase-amplitude coupling between oscillatory gamma and alpha activity, indicative of less modulated multisensory integration compared to non-falling older adults. As such, whilst analysing power within individual frequency bands is useful for identifying the functional role of specific types of neural oscillations, it is likely that with regards to multisensory integration, more holistic findings may come from analysing the synchronisation of multiple neural oscillations to understand how information from different senses is selected and bound together (Scurry et al., 2021).

### Functional ability predicted alpha power, but not audiovisual integration

The interaction between functional ability and age group was found to predict alpha power within the task, which provides strong support for the role of attentional control in the balance elements of functional ability. Not only is balance negatively affected by age-related challenges concerning musculoskeletal demands, medications causing dizziness and unisensory declines (Lim & Kong, 2022; Callis, 2016; Reed-Jones et al., 2013), but the weaker inhibitory abilities experienced by older adults is also a significant contributor to their increased risk of falls (Zhang et al., 2020).

Unpicking this significant interaction may lead to important insights into how younger and older adults employ attentional mechanisms for functional ability. That is, for younger adults, greater increases in alpha power from baseline (i.e. stronger inhibitory control) were associated with lower SPPB scores (i.e. weaker functional ability), which is surprising considering the role of attention and inhibition in the balance elements of functional ability. However, for older adults, greater increases in alpha power were associated with stronger functional ability (higher SPPB scores). Perhaps the reason for this difference lies in the age-related changes in the neural mechanisms relied upon for balance in each age group (Malcolm et al., 2021). Indeed, age-related declines in sensorimotor tracts within posture control loops result in the increased activation of cortical brain regions for balance maintenance in older adults (Pepper & Nuttall, 2023; Kahya et al., 2019; Malcolm et al., 2021; Ozdemir et al., 2018). For example, Ozdemir et al. (2018) found increased gamma activity in central, frontal and central-parietal areas of older adults when sensory information is compromised; these increases in gamma activity have previously been attributed to sustained attention (Slobounov et al., 2009). Ozdemir et al. (2018) postulated that older adults may allocate increased attentional resources to postural control than younger adults.

This is in line with the scaffolding theory of cognitive ageing (Reuter-Lorenz & Park, 2014; Oosterhuis et al., 2023), in which increased activation of frontal brain regions, associated with executive function, may be a compensatory strategy for older adults to maintain functional ability despite neural degeneration of balance centres (Oosterhuis et al., 2023; Kahya et al., 2019; Park & Reuter-Lorenz, 2009; Montero-Odasso et al., 2017). In the context of the current study, older adults displaying an association between increased alpha power and higher SPPB scores provides strong support for the role of inhibitory processes in functional ability, and perhaps reflects the increased involvement of cortical regions for supporting functional ability in older adults. In contrast, the negative relationship between alpha power and SPPB scores in younger adults may instead reflect the lesser role of inhibitory mechanisms for functional ability in this age group, whose sub-cortical sensorimotor tracts are intact, rendering balance a more automatic process. It is important that future research focusses on uncovering the age-related changes in the cortical mechanisms required for balance maintenance, as at the moment, the evidence into such changes appears to be limited (Malcolm et al., 2021; Ozedmir et al., 2018).

A potential reason as to why functional ability and balance ability did not predict audiovisual integration within the task (Model 1) could be that the older adults who participated in the study were very physically fit and able. This is evident in that the younger and older adults who participated in the current study displayed no significant differences in balance confidence (as measured by the Activities-Specific Balance Confidence scale) or in physical activity levels (as measured by the Rapid Assessment of Physical Activity). Whilst older adults may display increased audiovisual integration within the stream-bounce task, the high physical ability of these older adults may mask the effects that this less accurate integration has on their balance. As such, perhaps balance ability as measured in this study was not sensitive enough to predict age-related changes in audiovisual integration. Indeed, many clinical assessments of balance and fall risk appear to suffer from floor and ceiling effects and lack sensitivity to detect small changes in balance ability (Balasubmaranian, 2015; Rockwood et al., 2008; Yelnik & Bonan, 2008). The finding that participants’ balance ability did not predict audiovisual integration within this task may also be a promising indication that whilst older adults may experience increased audiovisual integration, regular exercise and maintaining strong physical wellbeing could reduce the effects that these maladaptive perceptual changes have on fall risk in older adults.

### Practical applications and future considerations

The roles of attention and inhibition in multisensory integration, and the weakening of cognitive abilities with healthy ageing, raises important questions regarding the treatments and therapies that could be designed to improve the integrative processes of older adults and reduce their risk of falls. That is, whilst strength and balance training has been proven to improve gait and thus potentially reduce fall risk in older adults during motor interventions (see Sherrington et al., 2008 for a detailed meta-analysis), the most effective programmes appear to come from combining physical and cognitive therapies (de Bruin et al., 2011; Pichierri et al., 2012; van het Reve & de Bruin, 2014), over a sustained period of time. For example, van het Reve & de Bruin (2014) implemented a combined motor and cognitive intervention with older adults, in which alongside an exercise programme, participants also received 12 weeks of cognitive training which included attending to task-relevant stimuli and suppressing task-irrelevant stimuli. The researchers found that after strength-balance-cognitive training, participants’ dual task costs during walking were significantly reduced and gait initiation was improved compared to participants who underwent strength-balance training alone. Taken together, perhaps combined physical and cognitive treatments could be effective in reducing the risk of falls in older adults (van het Reve & de Bruin, 2014; Uemura et al., 2012). However, when randomised control trials have been implemented amongst community-dwelling older adults, the findings have been mixed with regards to whether combined cognitive and physical interventions can reduce fall risk more than physical therapy in isolation (Turunen et al., 2022; Lipardo & Tsang, 2020; Segev-Jacubovski et al., 2011). As such, it is clear that further research is needed, with larger sample sizes and more diverse older adult populations, to determine whether such combined treatments are effective in minimising risk of falls in older adults.

The sampling bias that may be present in many studies investigating age-related changes in balance maintenance, or indeed any physical or cognitive aspect of ageing, must be taken into account in future research. For example, Brayne & Moffitt (2022) explained how ‘healthy volunteer bias’ is a high occurrence within ageing research, with older adults who agree to participate in such studies often being from more affluent subsections of society and healthier than randomly selected sample of the population. A consequence of this is that the results from studies using particularly healthy and able older adult samples may not be representative of the entire older adult population, making it difficult to generalise the findings (Brayne & Moffitt, 2022). However, it is important to note that these kinds of healthy volunteer biases are not necessarily limitations of ageing research, but instead, more detailed information about participants’ lifestyle, fitness, education and socialisation may be needed to create a more comprehensive account of the cognitive and physical abilities of the samples used – see Stern et al. (2020) and Oosterhuis et al. (2023) for reviews on the ‘cognitive reserve’ theories of ageing, which may contribute to the high level of individual differences within older adult groups.

## Conclusions

To conclude, the weaker top-down modulation of multisensory integration in older adults can have serious implications for their perception of and navigation through their dynamic environment. This study has provided support for the role of attentional control in functional ability, with age-related deteriorations in inhibitory function being a potential contributor to the increased risk of falls in older adults. To determine the underlying neural correlates of age-related changes in the top-down and bottom-up mechanisms of multisensory integration, and how these affect fall risk, it may be important to analyse neural activity from multiple frequency bands, to understand how oscillations coordinate to support multisensory perception and action. Future research must also investigate the possibility of younger and older adults using different strategies in facilitating the processing of task-relevant information and inhibiting task-irrelevant information; each age group may rely upon different brain areas and different mechanisms to support multisensory integration, compensating for age-related neurodegeneration. Specifically, the increased activation of cortical brain regions in older adults is likely to reflect their increased reliance on attentional and inhibitory mechanisms for balance maintenance, compared to younger adults. Developing a detailed understanding of the age-related changes in multisensory integration, and how this may influence fall risk, could provide important direction for the design of cognitive treatments to sharpen the perception of older adults and improve their allocation of attentional resources during balance maintenance.

## Appendix A Speech, Spatial and Quality of Hearing (SSQ) pre-screening questionnaire

**Start of Block: SSQ**

ssq info The following questions will ask you about aspects of your hearing ability, hearing experience and listening in different situations.

For each question, move the red circle along the slider and place it anywhere on the scale from 0 to 10.

Putting the red circle at 10 means that you would be perfectly able to do or experience what is described in the question. Putting the red circle at 0 means that you would be unable to do or experience what is described.

As an example, the first question asks about following a conversation with someone whilst the TV is on at the same time. If you are well able to do this, put the red circle at the right-hand end of the scale, at number 10. If you could follow about half the conversation in this situation, put the red circle around half-way along the scale, and so on.

Q9 You are talking with one other person and there is a TV on in the same room. Without turning the TV down, can you follow what the person you’re talking to says?

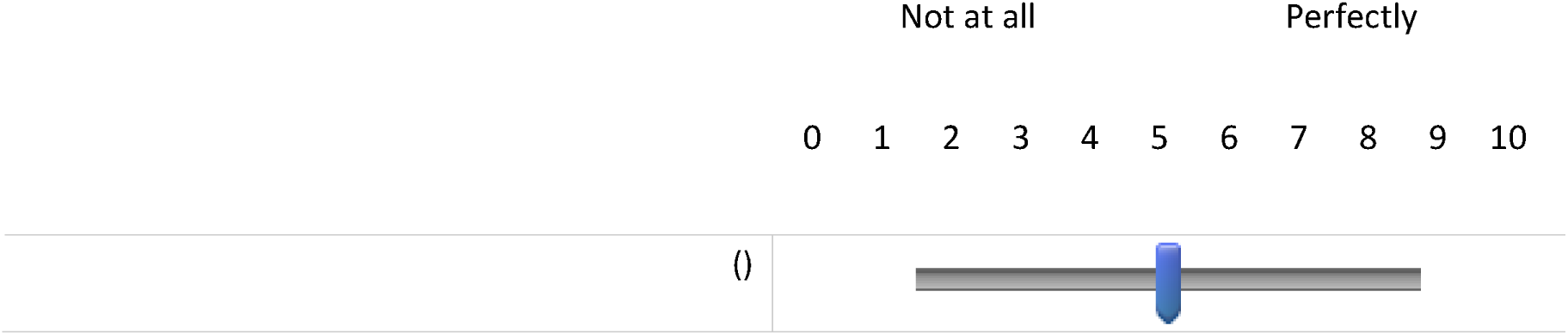

Q10 You are listening to someone talking to you, while at the same time trying to follow the news on TV. Can you follow what both people are saying?

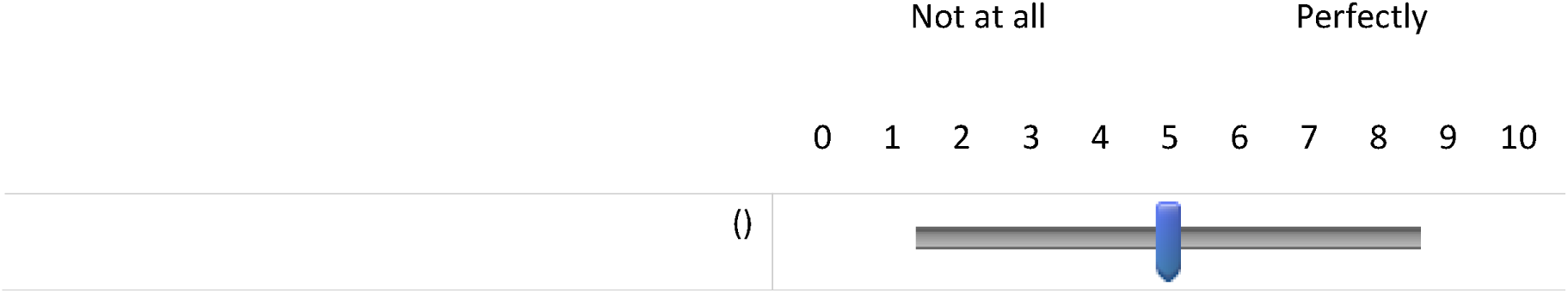

Q11 You are in conversation with one person in a room where there are many other people talking. Can you follow what the person you are talking to is saying?

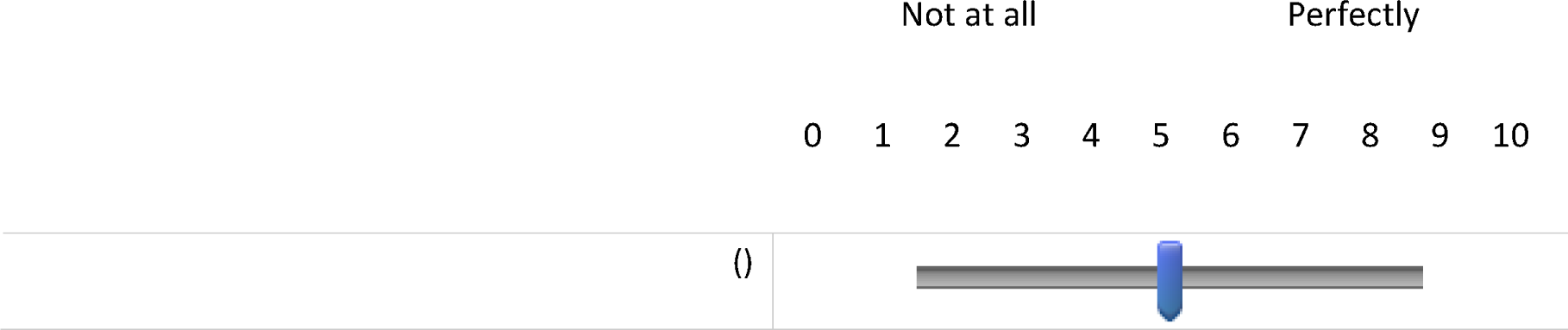

Q12 You are in a group of about five people in a busy restaurant. You can see everyone else in the group. Can you follow the conversation?

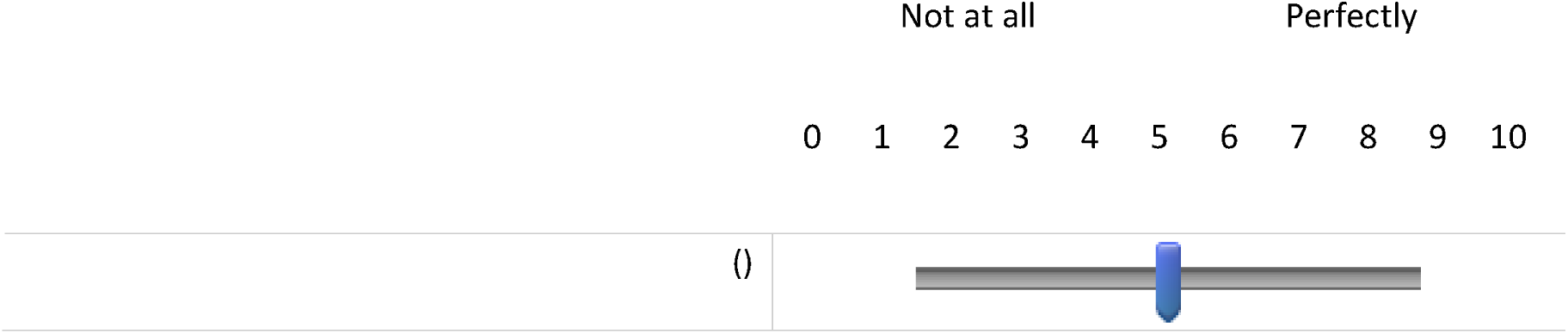

Q13 You are with a group and the conversation switches from one person to another. Can you easily follow the conversation without missing the start of what each new speaker is saying?

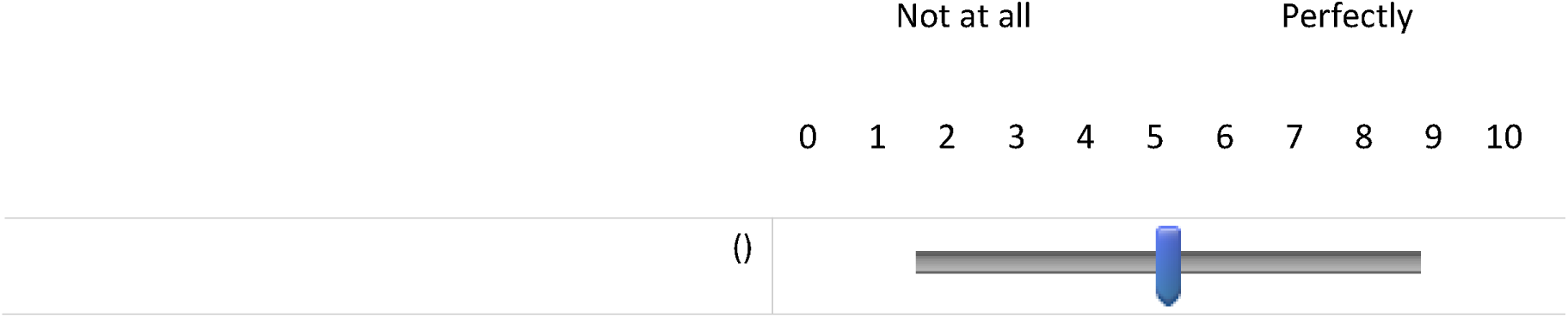

Q14 You are outside. A dog barks loudly. Can you tell immediately where it is, without having to look?

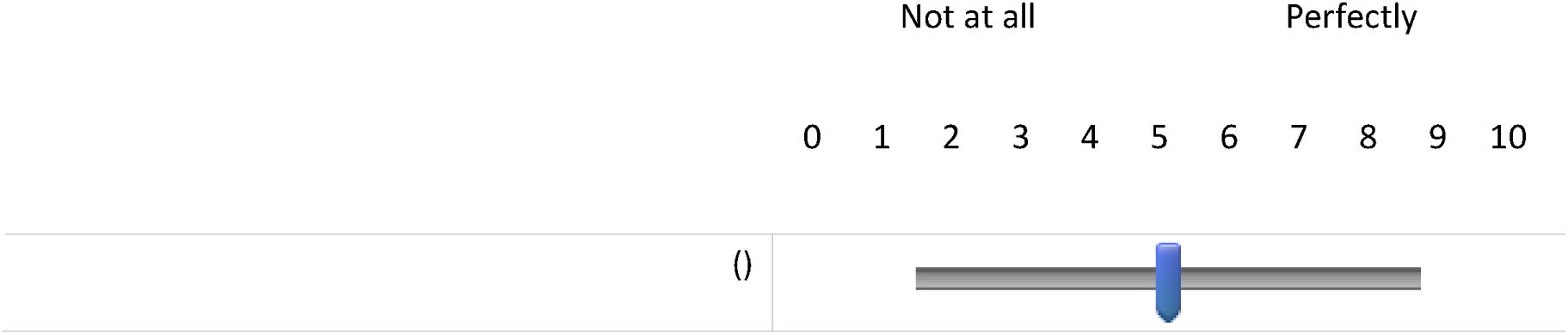

Q15 Can you tell how far away a bus or a truck is, from the sound?

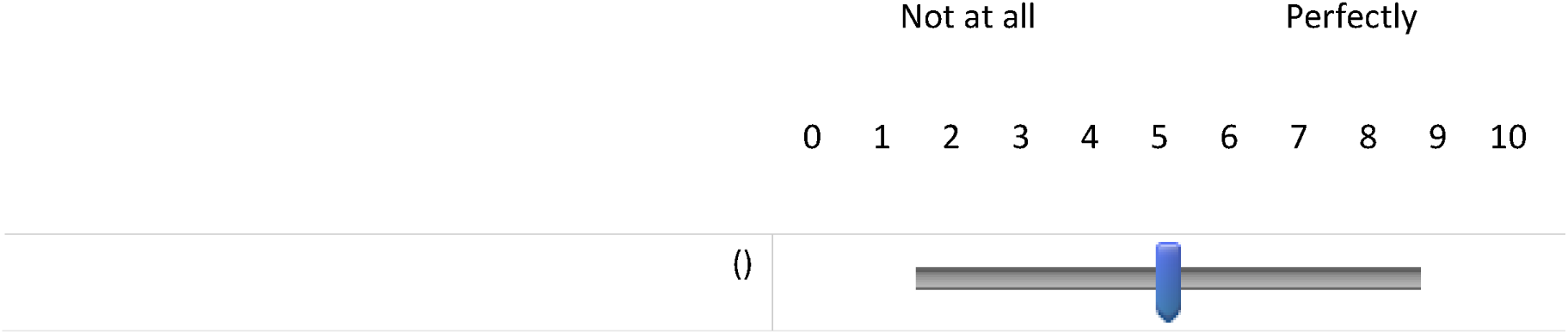

Q16 Can you tell from the sound whether a bus or truck is coming towards you or going away?

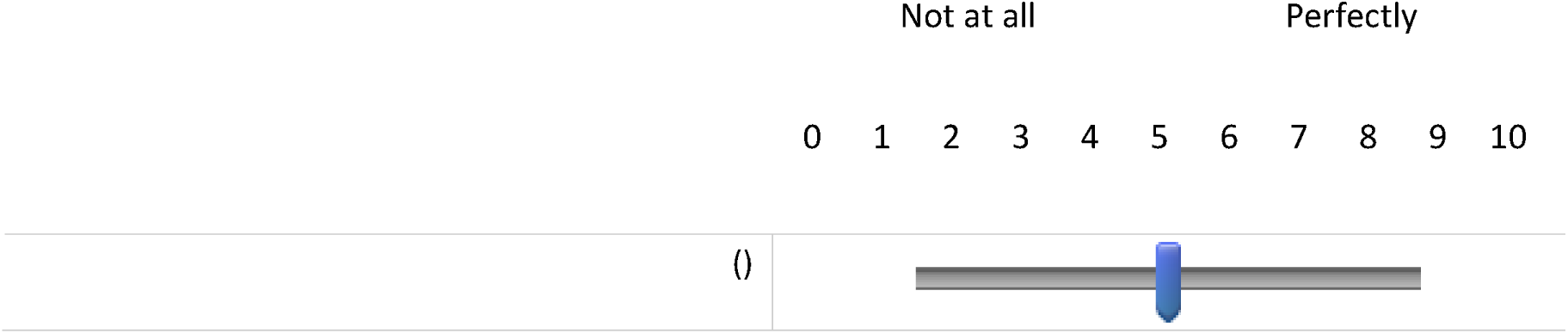

Q17 When you hear more than one sound at a time, do you have the impression that it seems like a single jumbled sound?

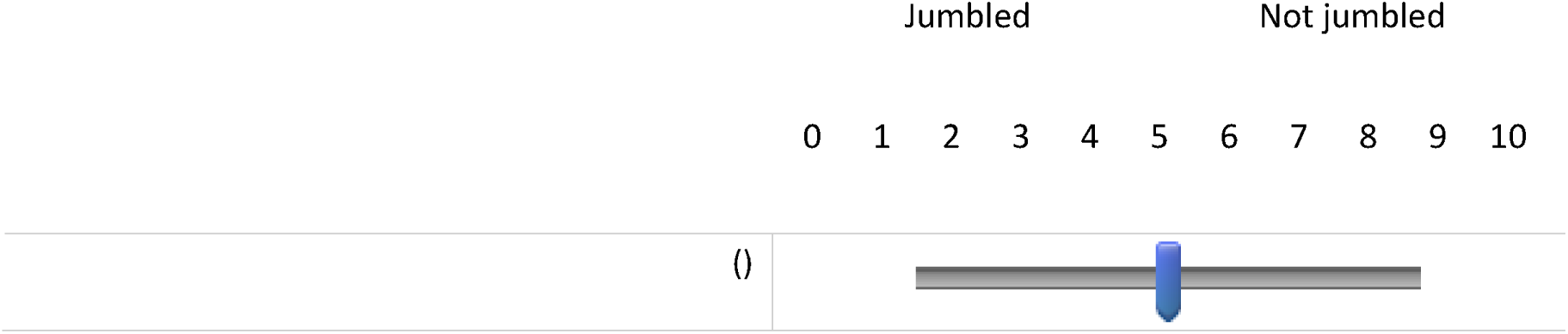

Q18 When you listen to music, can you make out which instruments are playing?

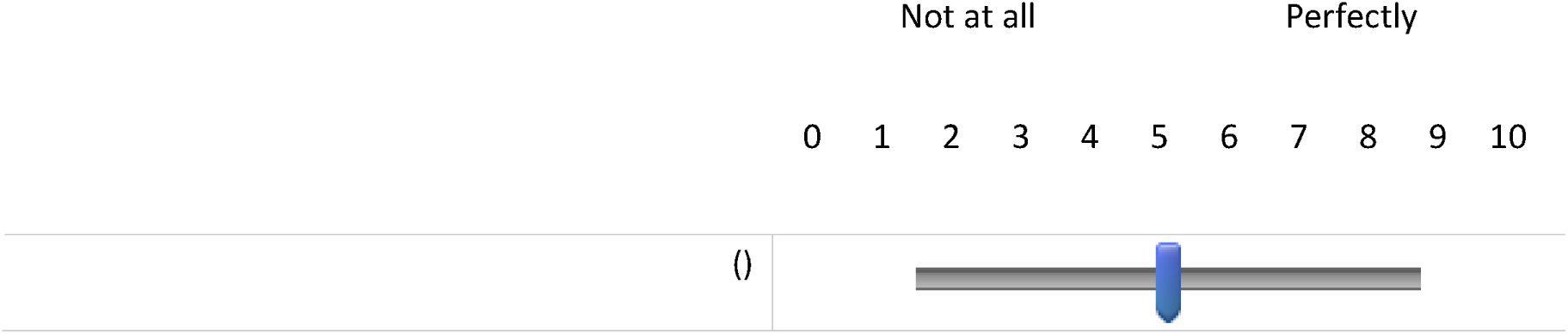

Q19 Do everyday sounds that you can hear easily seem clear to you (not blurred)?

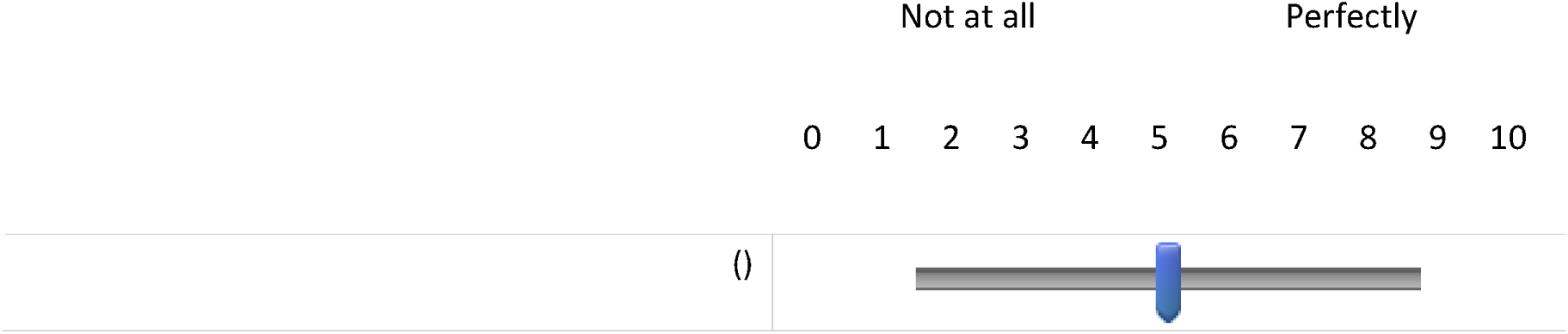

Q20 Do you have to concentrate very much when listening to someone or something?

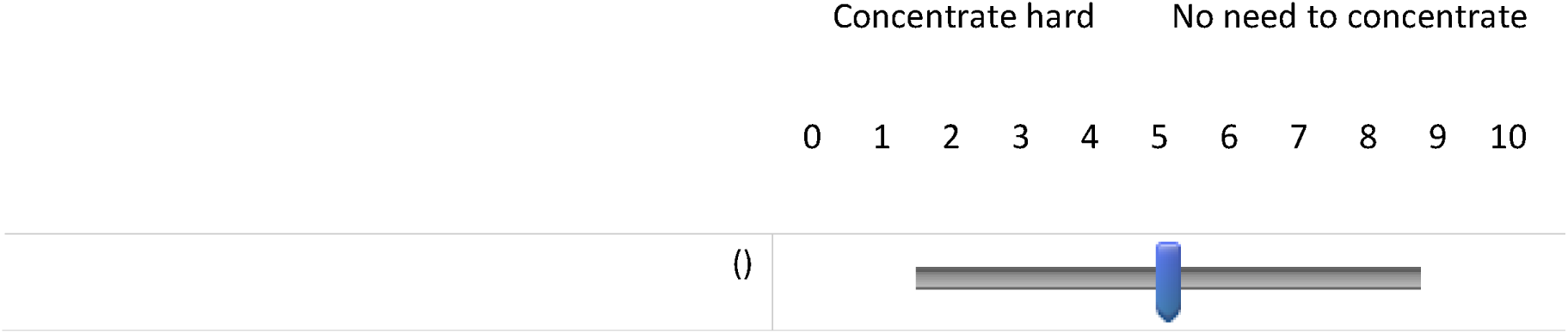

**End of Block: SSQ**

## Appendix B Informant Questionnaire on Cognitive Decline in the Elderly (IQ-CODE) pre-screening questionnaire

**Start of Block: IQCODE**

iqcode info Below are some situations where you have to use your memory or intelligence. Please indicate whether you have improved, stayed the same or got worse in that situation <u>**compared to 10** years</u>.

Q21 Compared to 10 years ago, how are you at remembering things about family and friends e.g. occupations, birthdays, addresses?

- Much improved (1)
- A bit improved (2)
- Not much change (3)
- A bit worse (4)
- Much worse (5)

Q22 Compared to 10 years ago, how are you at remembering things that have happened recently?

- Much improved (1)
- A bit improved (2)
- Not much change (3)
- A bit worse (4)
- Much worse (5)

Q23 Compared to 10 years ago, how are you at recalling conversations a few days later?

- Much improved (1)
- A bit improved (2)
- Not much change (3)
- A bit worse (4)
- Much worse (5)

Q24 Compared to 10 years ago, how are you at remembering your address and telephone number?

- Much improved (1)
- A bit improved (2)
- Not much change (3)
- A bit worse (4)
- Much worse (5)

Q25 Compared to 10 years ago, how are you at remembering what day and month it is?

- Much improved (1)
- A bit improved (2)
- Not much change (3)
- A bit worse (4)
- Much worse (5)

Q26 Compared to 10 years ago, how are you at remembering where things are usually kept?

- Much improved (1)
- A bit improved (2)
- Not much change (3)
- A bit worse (4)
- Much worse (5)

Q27 Compared to 10 years ago, how are you at remembering where to find things which have been put in a different place from usual?

- Much improved (1)
- A bit improved (2)
- Not much change (3)
- A bit worse (4)
- Much worse (5)

Q28 Compared to 10 years ago, how are you at knowing how to work familiar machines around the house?

- Much improved (1)
- A bit improved (2)
- Not much change (3)
- A bit worse (4)
- Much worse (5)

Q29 Compared to 10 years ago, how are you at learning to use a new gadget or machine around the house?

- Much improved (1)
- A bit improved (2)
- Not much change (3)
- A bit worse (4)
- Much worse (5)

Q30 Compared to 10 years ago, how are you at learning new things in general?

- Much improved (1)
- A bit improved (2)
- Not much change (3)
- A bit worse (4)
- Much worse (5)

Q31 Compared to 10 years ago, how are you at following a story in a book or on TV?

- Much improved (1)
- A bit improved (2)
- Not much change (3)
- A bit worse (4)
- Much worse (5)

Q32 Compared to 10 years ago, how are you at making decisions on everyday matters?

- Much improved (1)
- A bit improved (2)
- Not much change (3)
- A bit worse (4)
- Much worse (5)

Q33 Compared to 10 years ago, how are you at handling money for shopping?

- Much improved (1)
- A bit improved (2)
- Not much change (3)
- A bit worse (4)
- Much worse (5)

Q34 Compared to 10 years ago, how are you at handling financial matters e.g. the pension, dealing with the bank?

- Much improved (1)
- A bit improved (2)
- Not much change (3)
- A bit worse (4)
- Much worse (5)

Q35 Compared to 10 years ago, how are you at handling other everyday arithmetic problems e.g. knowing how much food to buy, knowing how long between visits from family or friends?

- Much improved (1)
- A bit improved (2)
- Not much change (3)
- A bit worse (4)
- Much worse (5)

Q36 Compared to 10 years ago, how are you at using your intelligence to understand what is going on and to reason things through?

- Much improved (1)
- A bit improved (2)
- Not much change (3)
- A bit worse (4)
- Much worse (5)

**End of Block: IQCODE**

## Appendix C Activities-Specific Balance Confidence scale (ABC)

Q1 Have you had a fall in the past?

- Yes (1)
- No (2)

*Display This Question:*

*If Have you had a fall in the past? = Yes*

Q2 How long ago was your fall? (Please answer in days, weeks, months or years)

Q5 For each of the following activities, please indicate your level of confidence in doing the activity, without losing your balance or becoming unsteady.

To do so, choose one of the percentage points on the scale from 0 (no confidence) to 10 (complete confidence)

If you do not currently do the activity in question, try and imagine how confident you would be if you had to do the activity.

If you normally use a walking aid to do the activity or hold onto someone, rate your confidence as it you were using these supports.

Q6 Walk around the house?

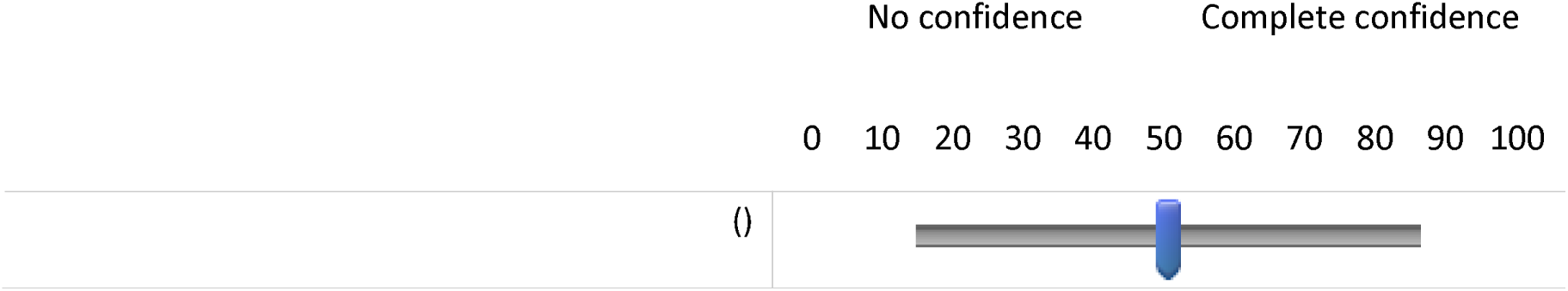

Q7 Walk up or down stairs?

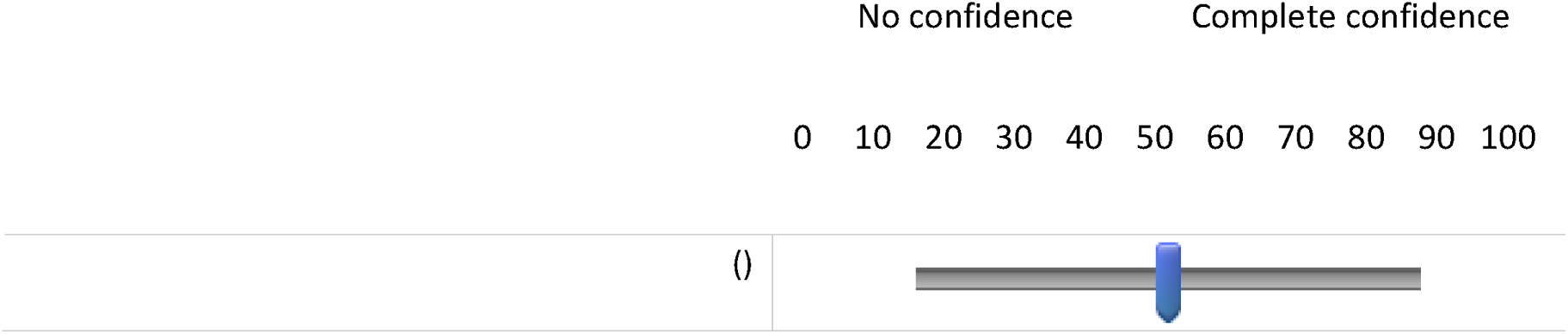

Q8 Bend over and pick up a slipper from the front of a closet floor?

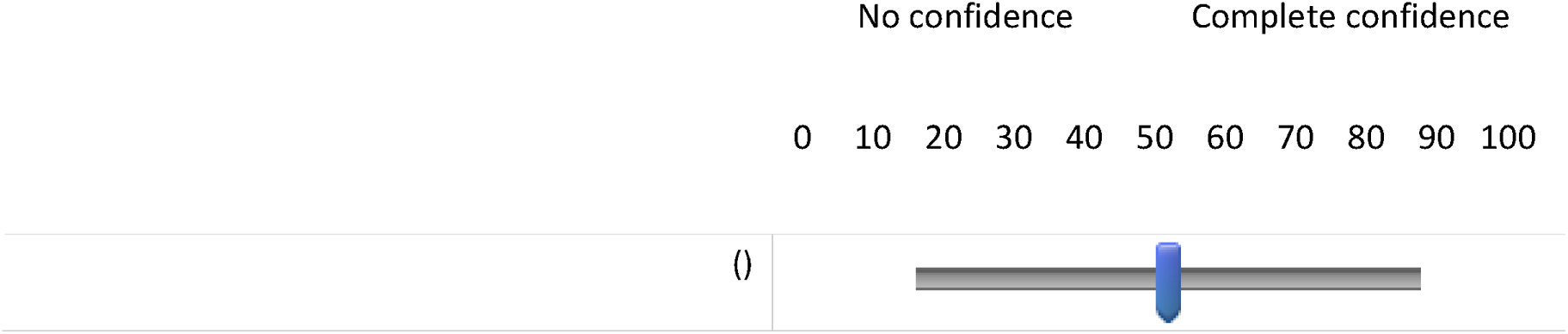

Q9 Reach for a small can off a shelf at eye level?

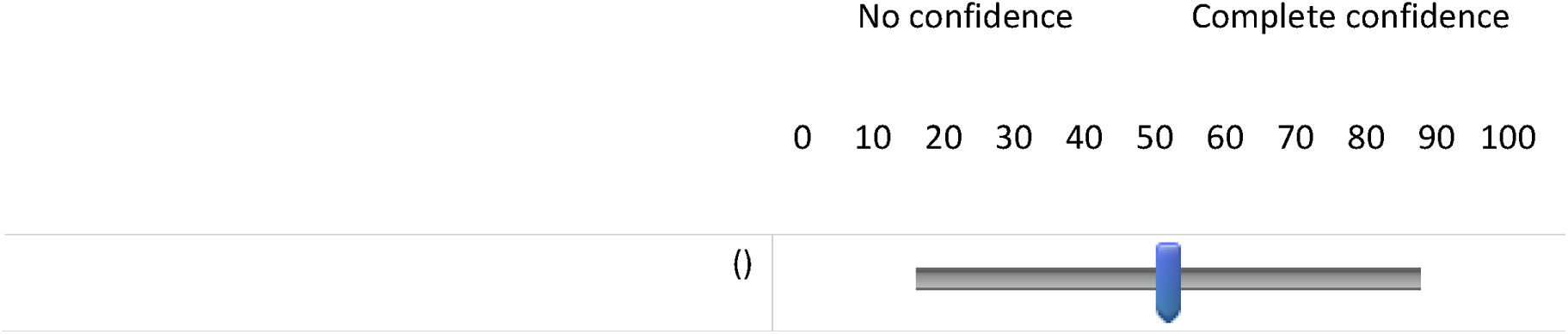

Q10 Stand on your tiptoes and reach for something above your head?

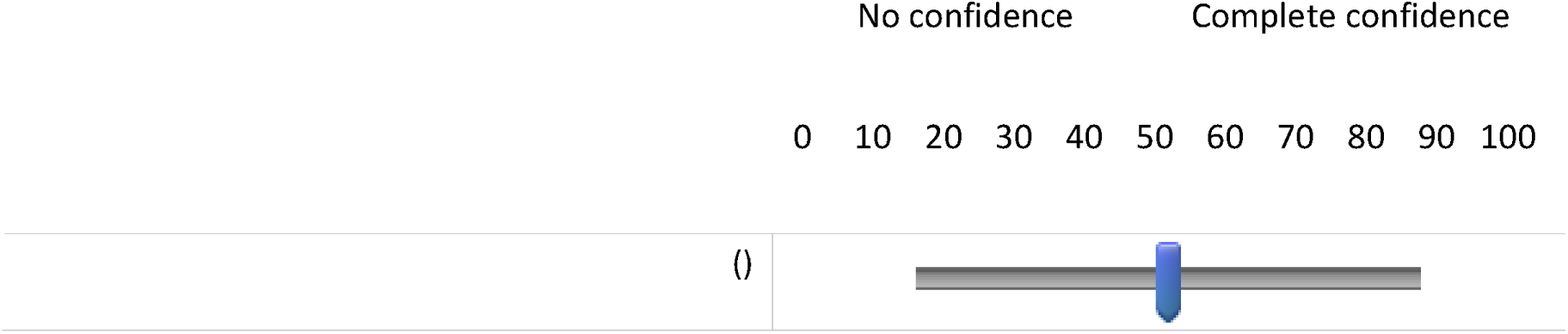

Q11 Stand on a chair and reach for something?

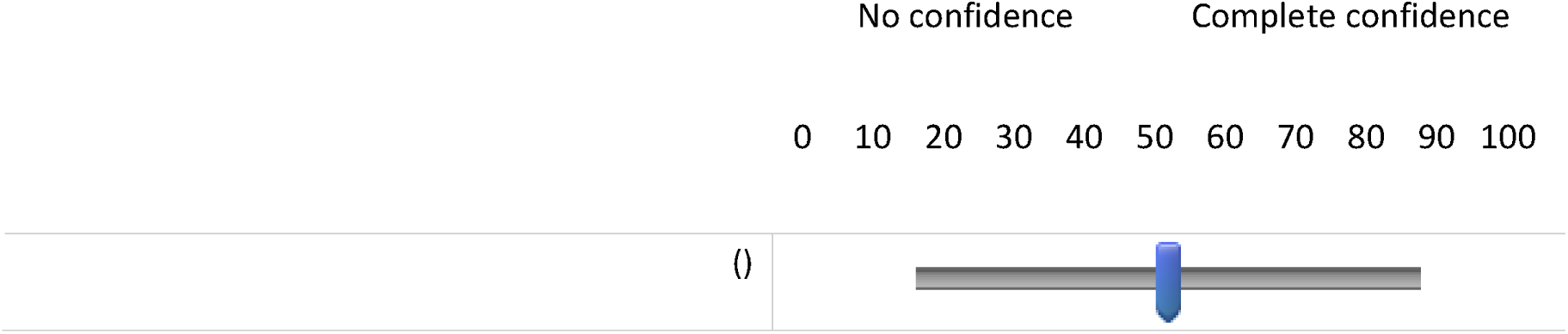

Q12 Sweep the floor?

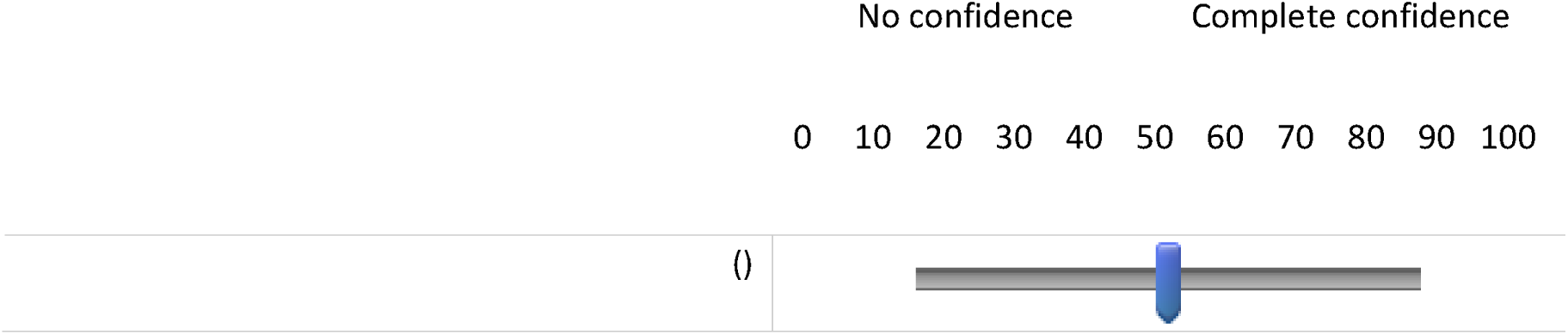

Q13 Walk outside the house to a car parked in the driveway?

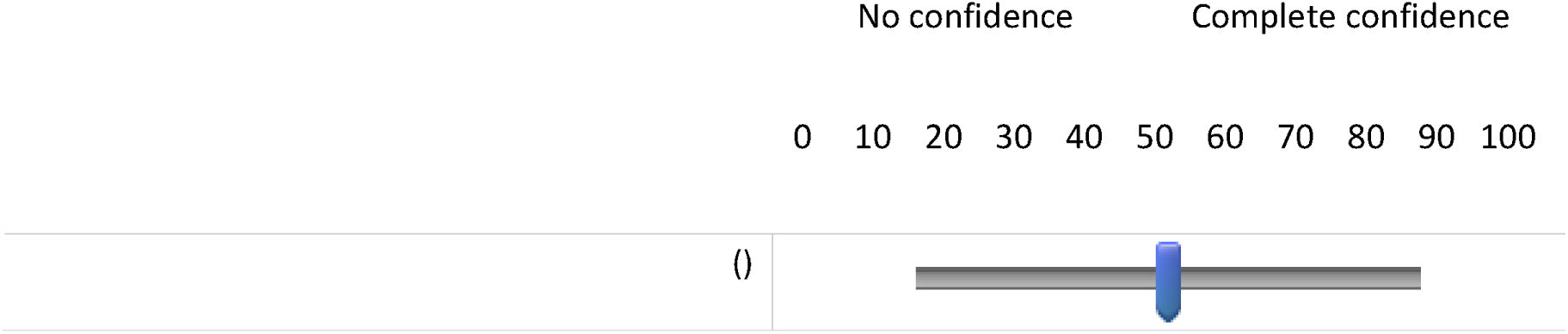

Q14 Get into or out of a car?

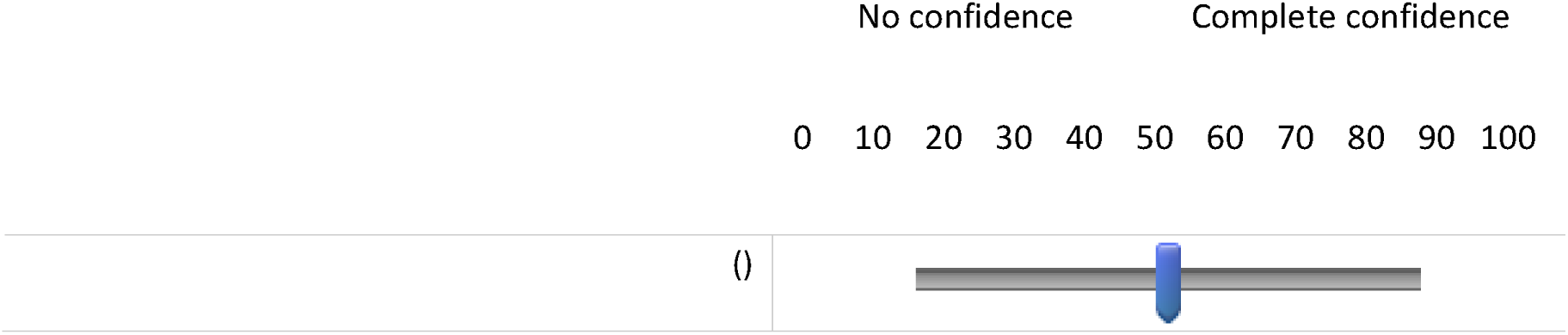

Q15 Walk across a car park to a shop?

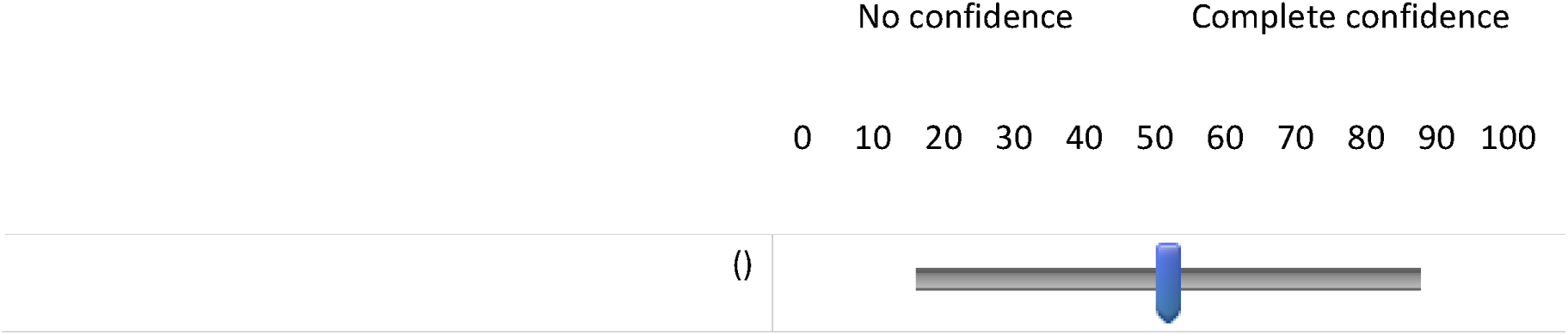

Q16 Walk up or down a ramp?

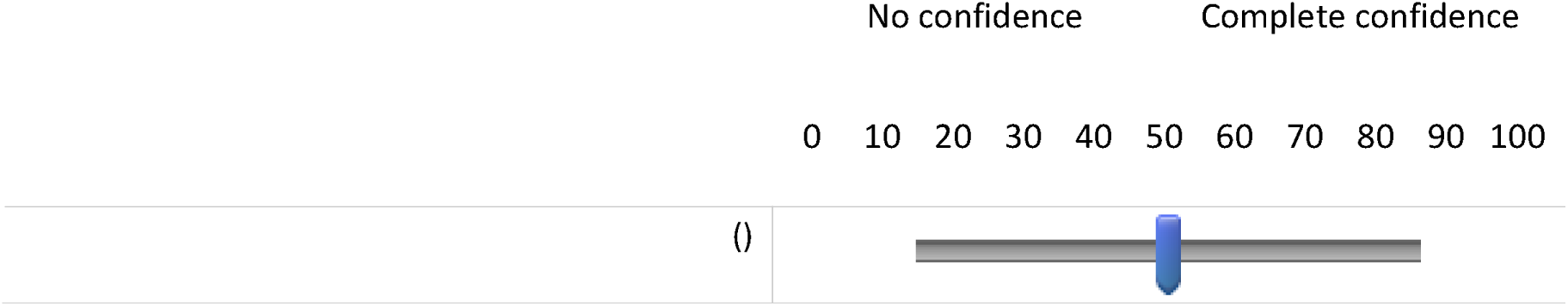

Q17 Walk in a crowded shop where people rapidly walk past you?

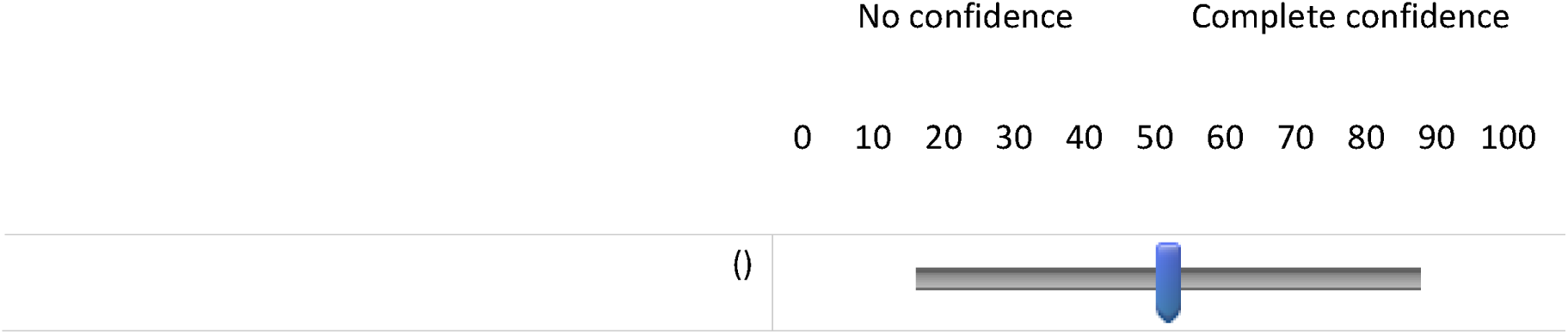

Q18 Are bumped into by people as you walk through the shop?

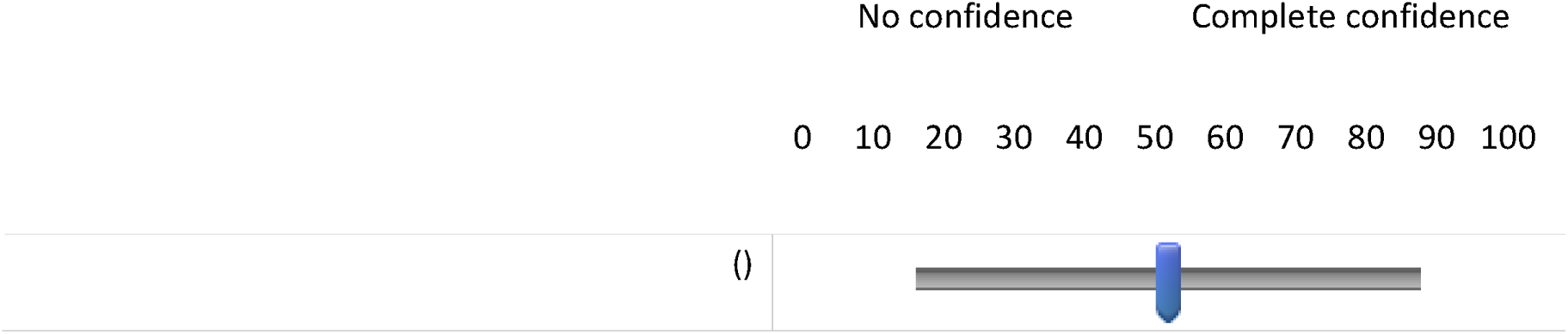

Q19 Step onto or off an escalator while you are holding onto a railing?

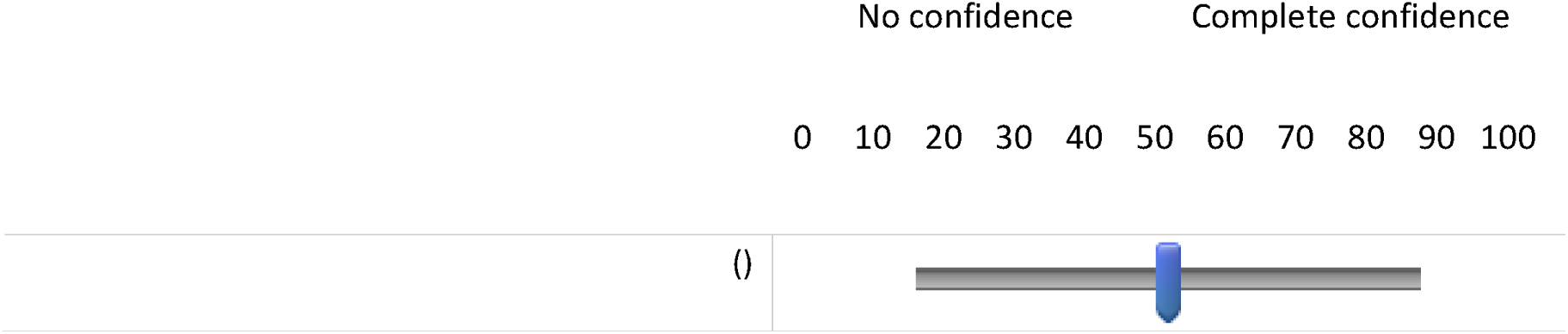

Q20 Step onto or off an escalator while holding onto parcels such that you cannot hold onto the railing?

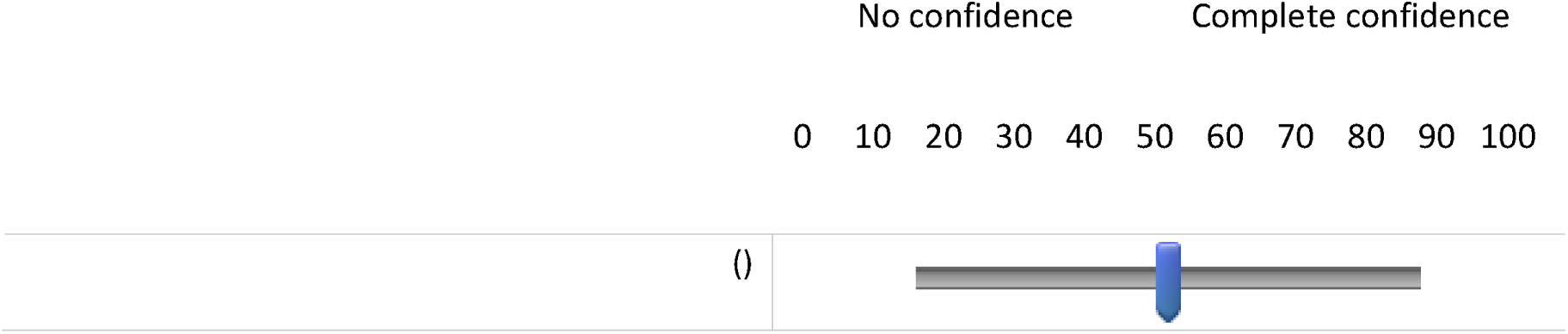

Q21 Walk outside on icy pavements?

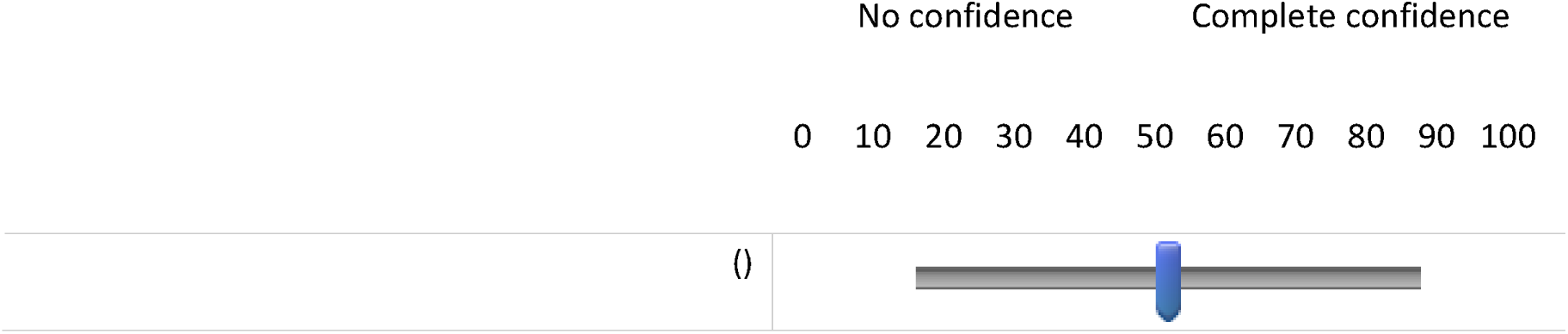

**End of Block: ABC Scale**

## Appendix D Rapid Assessment of Physical Activity (RAPA) questionnaire

**Start of Block: RAPA**

Q2 Physical Activities are activities where you move and increase your heart rate above its resting rate, whether you do them for pleasure, work, or transportation.

The following questions ask about the amount and intensity of physical activity you usually do. The intensity of the activity is related to the amount of energy you use to do these activities.

Light Activities:

- your heart beats slightly faster than normal
- you can talk and sing

Moderate Activities:

- your heart beats faster than normal
- you can talk but not sing

Vigorous Activities

- your heart rate increases a lot
- you can’t talk or your talking is broken up by large breaths

Q8 How physically active are you? Please check either "Yes" or "No" on each line whether you think that the statement accurately describes you.

Q9 I rarely or never do any physical activities.

- Yes (1)
- No (2)

Q10 I do some light or moderate physical activities, but not every week.

- Yes (1)
- No (2)

Q11 I do some light physical activity every week.

- Yes (1)
- No (2)

Q12 I do moderate physical activities every week, but less than 30 minutes a day or 5 days a week.

- Yes (1)
- No (2)

Q13 I do vigorous physical activities every week, but less than 20 minutes a day or 3 days a week.

- Yes (1)
- No (2)

Q14 I do 30 minutes or more a day of moderate physical activities, 5 or more days a week.

- Yes (1)
- No (2)

Q15 I do 20 minutes or more a day of vigorous physical activities, 3 or more days a week.

- Yes (1)
- No (2)

Q16 I do activities to increase muscle strength, such as lifting weights or calisthenics, once a week or more.

- Yes (1)
- No (2)

Q17 I do activities to improve flexibility, such as stretching or yoga, once a week or more.

- Yes (1)
- No (2)

**End of Block: RAPA**

